# Meiosis-specific prophase-like pathway controls cleavage-independent release of cohesin by Wapl phosphorylation

**DOI:** 10.1101/250589

**Authors:** Kiran Challa, V Ghanim Fajish, Miki Shinohara, Franz Klein, Susan M. Gasser, Akira Shinohara

## Abstract

Sister chromatid cohesion on chromosome arms is essential for the segregation of homologous chromosomes during meiosis I while it is dispensable for sister chromatid separation during mitosis. It was assumed that, unlike the situation in mitosis, chromosome arms retain cohesion prior to onset of anaphase-I. Paradoxically, reduced immunostaining signals of meiosis-specific cohesin, including the kleisin Rec8, from the chromosomes were observed during late prophase-I of budding yeast. This decrease is seen in the absence of Rec8 cleavage and depends on condensin-mediated recruitment of Polo-like kinase (PLK/Cdc5). In this study, we confirmed that this release indeed accompanies the dissociation of acetylated Smc3 as well as Rec8 from meiotic chromosomes during late prophase-I. This release requires, in addition to PLK, the cohesin regulator, Wapl (Rad61/Wpl1 in yeast), and Dbf4-dependent Cdc7 kinase (DDK). Meiosis-specific phosphorylation of Rad61/Wpl1 and Rec8 by PLK and DDK collaboratively promote this release. This process is similar to the vertebrate “prophase” pathway for cohesin release during G2 phase and pro-metaphase. In yeast, meiotic cohesin release coincides with PLK-dependent compaction of chromosomes in late meiotic prophase-I. We suggest that yeast uses this highly regulated cleavage-independent pathway to remove cohesin during late prophase-I to facilitate morphogenesis of condensed metaphase-I chromosomes.

**Author Summary:** In meiosis the life and health of future generations is decided upon. Any failure in chromosome segregation has a detrimental impact. Therefore, it is currently believed that the physical connections between homologous chromosomes are maintained by meiotic cohesin with exceptional stability. Indeed, it was shown that cohesive cohesin does not show an appreciable turnover during long periods in oocyte development. In this context, it was long assumed but not properly investigated, that the prophase pathway for cohesin release would be specific to mitotic cells and will be safely suppressed during meiosis so as not to endanger the valuable chromosome connections. However, a previous study on budding yeast meiosis suggests the presence of cleavage-independent pathway of cohesin release during late prophase-I. In the work presented here we confirmed that the prophase pathway is not suppressed during meiosis, at least in budding yeast and showed that this cleavage-independent release is regulated by meiosis-specific phosphorylation of two cohesin subunits, Rec8 and Rad61(Wapl) by two cell-cycle regulators, PLK and DDK. Our results suggest that late meiotic prophase-I actively controls cohesin dynamics on meiotic chromosomes for chromosome segregation.

## Introduction

Meiosis gives rise to haploid gametes from diploid germ cells. During meiosis, a single round of DNA replication is followed by two consecutive chromosome segregations, meiosis I and II, which reduce the number of chromosomes by half [1]. Homologous chromosomes are separated during meiosis I (MI), and sister chromatids are segregated during meiosis II (MII). Sister chromatid cohesion (SCC) acts as physical connection between the segregating chromosomes and provides resistance to pulling forces by microtubules. SCC along chromosome arms and at the kinetochore plays a critical role in chromosome segregation during MI and MII, respectively. For accurate chromosome segregation at MI, SCC along chromosome arms, and chiasmata, which are the cytological manifestation of crossovers, are essential for generating tension between the homologous chromosomes.

SCC is mediated by a protein complex, called cohesin, that is able to embrace two sister chromatids in a ring-shaped structure [2]. The core subunits of cohesin are composed of two structure-maintenance complex (SMC) ATPases, Smc1 and Smc3, as well as a kleisin subunit, Scc1/Mcd1/Rad21 (hereafter, Scc1 for simplicity). Smc1 and Smc3, both of which consist of a rod-like structure with an ATPase head, form a heterodimeric ring, which entraps two DNA duplexes. Scc1 bridges between the Smc1 and Smc3 ATPase head domains to lock the ring.

Chromosomal localization of cohesin is highly dynamic, and is strictly regulated. During the G1 phase, the loading of cohesin is mediated by the Scc2-Scc4 loader complex [3]. This process itself is not sufficient for SCC formation. However, SCC establishment occurs in S phase, during which the Eco1 acetyl-transferase catalyzes Smc3 acetylation [4-6]. SCC is thereafter maintained until the onset of anaphase, when Scc1 is cleaved by the protease separase [7]. This results in the release of the two entrapped sister chromatids. Separase activity is regulated by the protein securin, which binds to separase to inhibit its function. This process is closely monitored by the spindle-assembly checkpoint (SAC) to ensure that each chromosome is properly attached to the spindle apparatus prior to separation [1]. SAC negatively controls the protein ubiquitination machinery, the anaphase-promoting complex/cyclosome (APC/C), whose activation is essential for entry into anaphase. Activation of APC/C requires the Cdc20 and APC/C-Cdc20 targets securin for destruction, which in turn enables the separation of sister chromatids. Thus, the activity of Cdc20 plays a critical role in Scc1 cleavage, and consequently, the transition from metaphase to anaphase.

Cohesin dynamics are regulated by other cohesin-interacting proteins in yeasts and vertebrates, such as Scc3, Pds5 and Rad61/Wpl1 (Wapl), with vertebrates also having a Wapl antagonist called sororin [8]. Wapl, together with Pds5, negatively regulates the binding of cohesin to chromatin [9, 10]. Wapl-regulated cohesin dissociation is independent of Scc1 cleavage, allowing entrapped DNAs to be released from the cohesin ring by opening the “exit gate” at the interphase between Smc3 and Scc1 [11, 12]. Eco1-mediated Smc3 acetylation locks the gate [5, 13] and sororin interacts with the cohesin complex to suppress Wapl activity [8].

In vertebrate cells during late G2 or pro-metaphase, cohesin is removed from the majority of chromosome arms by a Scc1-cleavage-independent pathway [14]. This so-called “prophase pathway” for cohesin removal is triggered by the phosphorylation of sororin and Scc3 by polo-like kinase (PLK), aurora kinase, and cyclin-dependent kinase (CDK) [8, 15]. Phosphorylated sororin is inactive, and can no longer suppress Wapl activity. On the other hand, at kinetochores, the phosphorylation that triggers the prophase pathway is blocked by the action of Shugoshin, a protein that recruits a phosphatase, PP2A [16]. PP2A is believed to dephosphoryate proteins involved in the prophase pathway, such as sororin. Interestingly, sororin is not present in lower eukaryotes such as budding yeast, and the prophase pathway of cohesin removal is absent in yeast mitosis [17].

Cohesin also plays an essential role in chromosome segregation during meiosis [1]. During meiosis, the kleisin Scc1 is replaced with its meiosis-specific counterpart, Rec8 (and also RAD21L in mammals) [18-20]. The Rec8-cohesin complex is also involved in various chromosomal events such as homologous recombination and chromosome motion in the meiotic prophase-I [21, 22]. Cohesin is a major component of the chromosome axis, which contains two sister chromatids organized into multiple chromatin loops [19]. During the pachytene stage, homologous chromosomes pair with each other, and synapse along chromosome axes to form a unique meiosis-specific chromosome structure, the synaptonemal complex (SC) [23]. The SC together with chromosome axes then dismantle to form chiasmata during diplonema and diakinesis/early metaphase-I. At the onset of anaphase-I, APC/C-Cdc20 induces securin degradation, which activates separase to allow cleavage of Rec8. Phosphorylation of Rec8 by three kinases, PLK, Dbf4-dependent Cdc7 kinase (DDK), and Casein kinase 1 (CK1) promotes this cleavage [24, 25]. Upon the onset of anaphase-I, the phosphorylation and cleavage of Rec8 is restricted to chromosome arms, while Rec8 at the kinetochores is protected by Shugoshin [26]. It has been shown that Shugoshin blocks phosphorylation of Rec8 at the kinetochores by recruiting PP2A [27, 28]. Protection of SCC at the kinetochores is essential for proper sister chromatids segregation at MII.

Previously, Yu and Koshland (2005) analyzed the role of condensin, a related SMC complex that is required for chromosome condensation in mitosis, for the resolution of recombination-dependent linkage between homologous chromosomes. Their immuno-staining showed a decreased intensity of Rec8-cohesin signal on meiotic chromosomes during late prophase I relative to mid-prophase I. The decreased Rec8 signals were also seen in a separase mutant defective in Rec8-cleavage (*esp1-1*) as well as a cell arrested prior to anaphase I. These results suggested that a subset of Rec8-cohesin is released from meiotic chromosomes during late prophase I, as in vertebrate mitosis. Importantly, this Rec8 release required the condensin-dependent recruitment of Cdc5/PLK to the chromosomes. However, the Cdc5 target involved in Rec8-release during meiosis remained unidentified, even though the study suggested a potential role for Rec8 phosphorylation in the pathway. Their proposed role for Cdc5/PLK in Rec8 dynamics remained a bit controversial [24, 25, 29]. A recent study in *C. elegans* indicated a role for condensin in the “retention” of meiotic cohesin complexes on meiotic chromosomes at least during early- and mid-prophase I by antagonzing Wapl activity [30].

Here, we revisited the question of Rec8-cohesin dynamics during late prophase-I and confirmed that, in late prophase-I of budding yeast, approximately half of the full-length of Rec8 molecules are released from meiotic chromosomes in a cleavage-independent manner [31]. In addition to Cdc5/PLK, we showed that this “prophase-like” removal of cohesin during meiotic prophase-I requires Rad61/Wpl1 and DDK. Furthermore, we found that not only DDK- and PLK-mediated phosphorylation of Rec8, but also meiosis-specific phosphorylation of Rad61/Wpl1 promotes Rec8-cohesin release. The Rec8 release is coupled to changes in chromosome compaction. We propose that cleavage-independent release of cohesin is a key regulator of meiotic chromosome function in late prophase-I.

## Results

### Rec8 shows dynamic localization during late prophase-I

Here we studied the dynamics of axis proteins during late prophase-I. Because late-prophase I is a very short-lived stage in budding yeast meiosis (e.g. Fig. 1G; ∼1.5 h from mid-pachytene and post-meiosis I in wild-type cells), we arrested cells prior to the onset of anaphase I using a meiosis-specific depletion mutant of *CDC20, cdc20mn* (meiotic null), which compromises activation of the anaphase promoting complex/cyclosome (APC/C) [32]. We then analyzed the localization of chromosome axis proteins in this *cdc20mn* mutant over a meiotic time course. The staining of a central component of the SC, Zip1 [33], allowed us to classify stages of prophase-I as follows: I, dotty staining; II, short-line staining; III, long-line staining. Category III corresponds to pachytene stage, during which chromosome synapsis occurs. Following pachytene stage, the SC dismantles, resulting in the re-appearance of dotty Zip1 staining (class-I), some of which co-localizes with kinetochores (see below) [34]. Disassembly of Zip1 was also found to correlate with dissociation of chromosome axis proteins such as Red1 (Fig 1A and 1B)[35].

**Figure 1.**
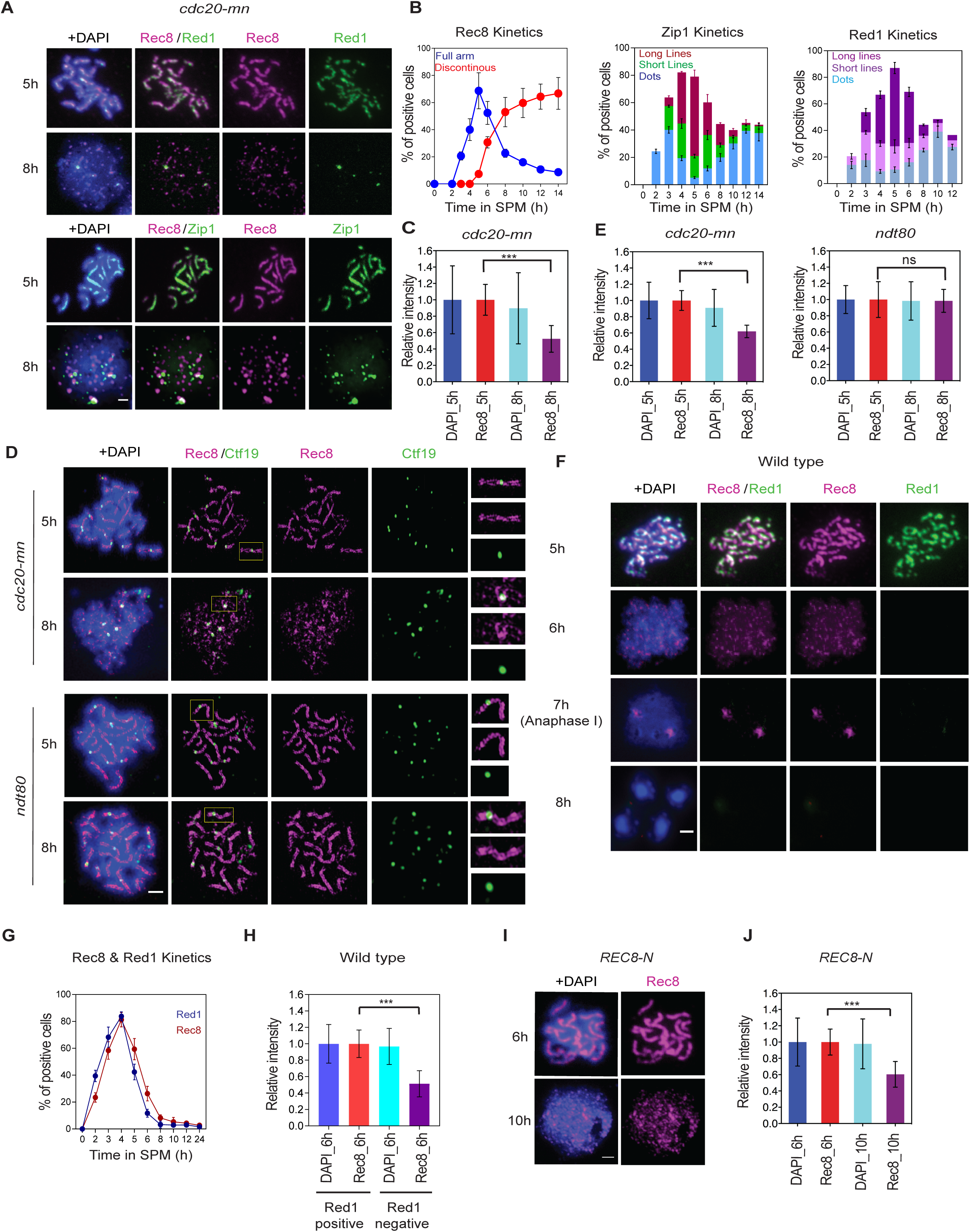
Rec8 shows dynamic localization in late meiotic prophase-I. **A**. Immunostaining analysis of Rec8 (red) and axis protein Red1 (green; top) and Rec8 (red) and SC protein Zip1 (green; bottom) in *cdc20-mn* (KSY642/643) strain. Representative image with or without DAPI (blue) dye is shown. Rec8 staining in the *cdc20-mn* was classified as linear (5 h) and altered (8 h) classes. The bar indicates 2 µm. **B** Kinetics of Rec8 (left), Zip1 (middle) and Red1 (right) staining in *cdc20-mn* (KSY642/643) strain was analyzed. A minimum 100 cells were counted at each time point. Error bars (Rec8) show the standard deviation (S.D.; n=3). Rec8 staining is classified; full (blue) and discontinuous dotty (red) staining. Zip1 staining is classified as follows: dotty (blue); short linear (green); full linear staining (red). Red1 staining is classified as follows; dotty (light blue); short linear (light purple); full linear staining (purple). **C**. Total signal intensity of Rec8 and DAPI on chromosome spreads at 5 and 8 h was measured in *cdc20-mn* (KSY642/643) cells. A minimum 30 spreads were quantified in representative time points. Error bars show the S.D. (n=3). **D**. SR-SIM microscopic observation of Rec8 (red) and Ctf19 (green) (left) in *cdc20-mn* (KSY642/643) and *ndt80* (KSY467/468) cells. Representative image with or without DAPI (blue) dye is shown. White insets are shown in a magnified view at right. The bar indicates 2µm.**E**. Total signal intensity of Rec8 and DAPI on chromosome spreads at 5 and 8 h was measured in *cdc20-mn* (KSY642/643) and *ndt80* (KSY467/468) cells as shown in (C). **F**. Immunostaining analysis of Rec8 (red) and Red1 (green) in wild type (MSY832/833) cells. Representative image with or without DAPI (blue) dye is shown. Rec8 staining was classified as linear (5 h) and altered (6 h) classes with Red1-positive and -negative, respectively. The bar indicates 2 µm. **G**. Kinetics of Rec8- and Red1-positive spreads in wild type (MSY832/833) is shown. A minimum 100 nuclei were quantified in representative time points. Error bars show the S.D. (n=3). **H**. Quantification of total Rec8 and DAPI signal intensity in Red1-postive and Red1-negative spreads in wild type was analyzed as shown in (C). **I**. Localization of Rec8 (red) with or without DAPI (Blue) in *REC8-N* (KSY597) mutants was analyzed and the representative images are shown. **J**. Quantification of total Rec8 and DAPI signal intensity at 5 and 8 h in *REC8-N* (KSY597) was analyzed as shown in (C).

We observed long lines of Zip1 staining peaks at 5 h in *cdc20mn* mutants, followed by the appearance of Zip1 dots after 6 h (Fig 1A and 1B). When localization of Rec8 was analyzed in pachytene stage, Rec8 shows linear staining that co-localized with Zip1-lines (Fig 1A). Following the pachytene stage, chromosome spreads with Zip1 dots (i.e. at 6 h) started to exhibit altered Rec8 staining associated with discontinuous dots (Fig 1A). This indicated that the remodeling of cohesin localization takes place at or after SC disassembly. Discontinuous dots of Rec8 staining in the *cdc20mn* mutants accumulated for up to ∼8 h. The appearance of discontinuous Rec8 staining occurred concomitantly with disassembly of SCs into Zip1-dots (Fig 1B). To further characterize the disassembly status of the axes, we also examined Rec8-Red1 localization at later time points; chromosomal Red1 signals diminish with a conversion from long lines to short lines/dots when discontinuous Rec8 staining is observed (Fig 1A and 1B). This result further confirmed that Rec8 remodeling occurs at or after disassembly of chromosome axis.

We then used super-resolution microscopy to analyze Rec8-cohesin localization on meiotic chromosomes at high resolution. A structural illumination microscope (SIM) was used to determine Rec8 localization in *cdc20mn* and *ndt80*, which arrests at the pachytene stage [36](Fig 1D). At 5 h, both strains showed two parallel lines of Rec8, which corresponded to two axes of full length SCs (Fig 1D). More importantly, by SIM the two Rec8 lines do not show a uniform staining, but rather a beads-on-string-like staining was observed. This suggested non-continuous localization of Rec8-cohesin along the chromosome axis. Kinetochore visualization by staining of Ctf19, a centromere protein, showed that each SC harbored a single focus of Ctf19, indicating tight fusion of all four sister kinetochores. Two linear Rec8 patterns were fused at the Ctf19 focal point, suggesting a unique axial structure at peri-centromeric regions with respect to Rec8-cohesin localization. In the *ndt80* mutant, these two Rec8 lines are maintained beyond 6 h. On the other hand, differential staining patterns were observed between 5 h and 8 h in *cdc20mn* mutants; the two clear parallel lines visible at 5 h disappear at 8 h, at which point discrete focus or short-line staining of Rec8 dominate. This is consistent with the observations made by conventional fluorescent microscopy described above. At later time points in *cdc20mn* mutants, a Ctf19 focus at kinetochores is often flanked by two distinct Rec8 signals (Fig 1D, see inset).

A previous study reported that the signal intensity of HA-tagged Rec8 was diminished during late prophase-I in *cdc20mn* as well as in wild type cells [31]. However, they did not observe the discontinuous Rec8 staining at late stages and instead a uniform staining of reduced intensity was seen. This may be due to differences in the antibodies used. In the previous study, the localization of HA-tagged Rec8 was examined using an anti-HA antibody. In our study, localization of non-tagged Rec8 was examined using two independent anti-Rec8 antisera. We quantified Rec8 signals on chromosomal spreads, and also found reduced Rec8 signal at 8 h (61.9±9.8%) as compared with that at 5 h in *cdc20mn* mutants (Fig 1E). This confirmed previous observations of HA-Rec8 [31]. This was also supported by quantification of Rec8 signal in our SIM images (Fig 1E). While epitope masking might also explain the decreased Rec8 signal, our results support the previous suggestion that Rec8 dissociates from chromosomes during late prophase-I [31]. Given that the decrease in Rec8 intensity was observed in *CDC20* depletion cells arrested at the meta/anaphase transition, Rec8 remodeling appeared to be independent of separase-mediated cleavage.

Because *CDC20* depletion may affect Rec8 localization during prolonged arrest, we confirmed Rec8 remodeling in late prophase-I by performing Rec8 and Red1 co-staining in a wild-type meiosis (Fig 1F and 1G). We examined spreads at 5 and 6 h when ∼70% cells are still in prophase-I, classifying these as Red1-positive and Red1-negative spreads, which correspond with pachytene/zygotene stages and diplotene/prometaphase-I, respectively. We then checked the staining pattern of Rec8 and its intensity. As seen upon *CDC20* depletion, the Red1-positive spreads showed linear Rec8 staining patterns (Fig 1F). On the other hand, Red1-negative spreads contained discontinuous dots of Rec8. This staining was different from that in anaphase-I spreads, which show two separated Rec8 foci with centromere clustering (Fig 1F, 7 h). Intensity measurements confirmed that Red1-negative spreads had reduced Rec8 signal (52.7±11.9% [n=20]) compared to Red1-positive spreads (100±15.1%; Fig 1H). Thus, Rec8 remodeling occurs during late prophase-I in normal wild-type meiosis as well.

To make sure that Rec8 depletion is independent of its cleavage, the localization of a Rec8 mutant protein, Rec8-N, which is resistant to cleavage by separase, was also investigated [37]. The *REC8-N* mutant strain shows normal prophase I progression, but is completely blocked at the metaphase/anaphase-I transition because Rec8-N is resistant to separase cleavage [37]. Similar to the *cdc20mn* mutant, at late prophase-I the *REC8-N* mutant exhibited discontinuous Rec8 staining with reduced intensity (60±15.7%), while linear staining was observed at 5 h (Fig 1I and 1J). Consistent with this, a previous study reported that the temperature-sensitive separase-deficient mutant, *esp1-1*, also showed a decrease in Rec8 intensity at late prophase-I at a restricted temperature [31]. Together, all observations support the hypothesis that Rec8 localization is remodeled and possibly released in late meiotic prophase-I in a manner independent of Rec8 cleavage.

### Rec8 dissociates from meiotic chromosomes at late prophase-I

To check whether Rec8-cohesin is indeed released from meiotic chromosomes in *cdc20mn* mutants, we fractionated cell/nuclear lysates to separate the chromatin bound and unbound fractions. Chromatin bound fractions contained proteins tightly bound to chromosomes, such as histones. At 5 h, most full-length Rec8 protein was recovered in the chromatin-bound fraction containing histone H2B in both *ndt80* and *cdc20mn* mutants (Fig 2A and S1A Fig). This argues that Rec8, and thus the cohesin complex, is tightly bound to DNA and/or chromatin. Rec8 is also chromatin-bound at 8 h in *ndt80* cells. On the other hand, a large proportion of Rec8 protein (66.0±15.7%) was recovered in the soluble fraction at 8 h in the *cdc20mn* mutant, and the remaining Rec8 protein was in the chromatin fraction (Fig 2A and 2B). Interestingly, the unbound Rec8 migrated more slowly on the gel than the bound Rec8. Rec8 is a target of Dbf4-dependent Cdc7 kinase (DDK), Polo-like kinase (Cdc5), and casein kinase 1 (Hrr25), and its slow-migrating form is highly phosphorylated [24, 25]. Its phosphorylation is believed to promote Rec8 cleavage by the separase, Esp1 [24, 25]. Importantly, Yu and Koshland (2005) showed that cohesin release is connected with Cdc5- and condensin-dependent phosphorylation of Rec8. Our results suggested that Rec8 phosphorylation triggers cohesin dissociation at late prophase-I.

**Figure 2.**
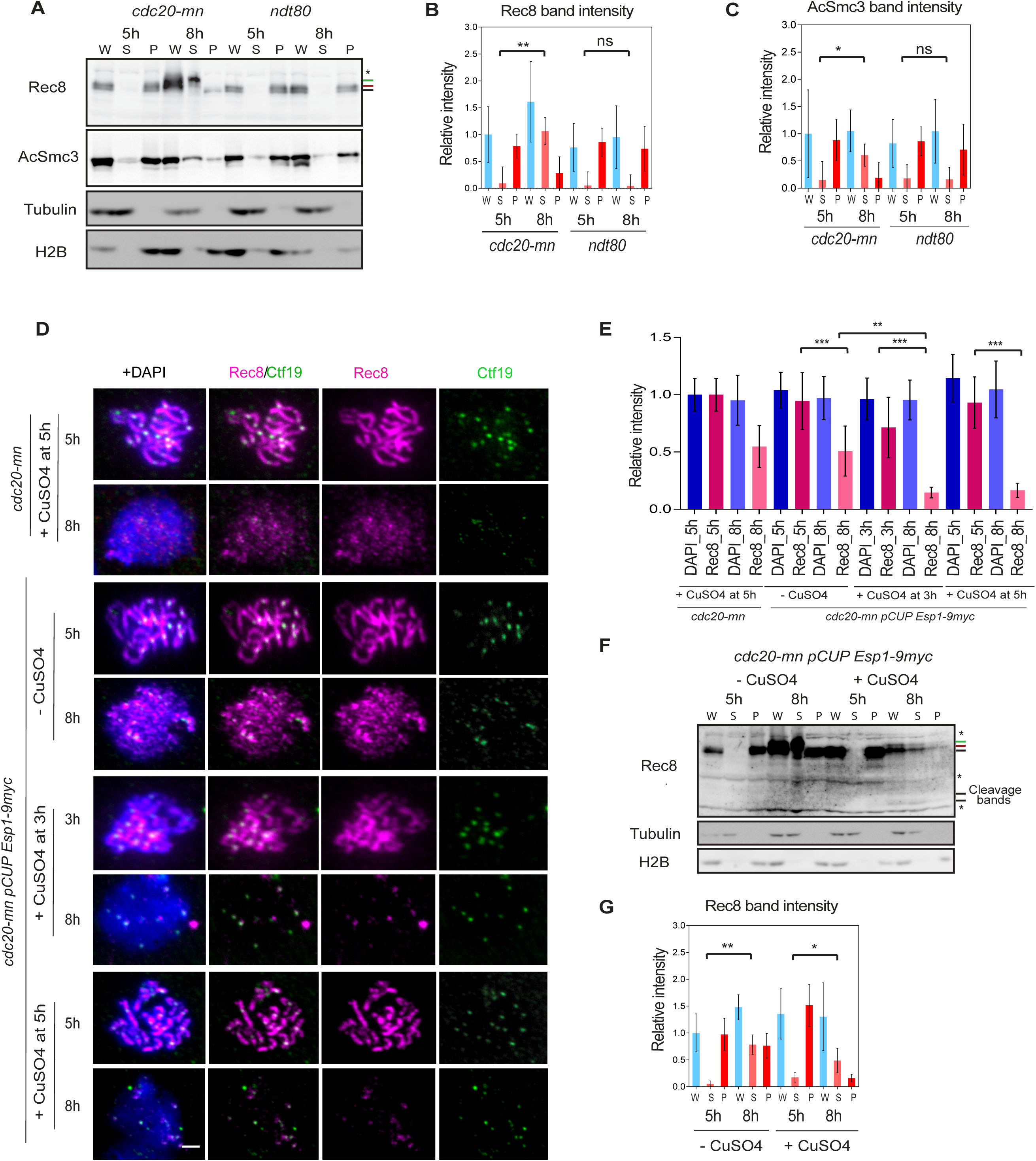
Rec8 dissociates from meiotic chromosomes at late prophase-I. **A**. Chromatin fractionation assay was carried out using *cdc20-mn* (KSY642/643) and *ndt80* (KSY467/468) mutant cells. Western blotting was performed for whole cell extracts (W), chromatin-unbound fractions (S) and chromatin-bound fraction (P). Rec8 (top) and acetyl-Smc3 (second) were probed together with tubulin (third) and Histone 2B (H2B; bottom) as controls for chromatin-unbound and -bound proteins, respectively. Two major phosphorylated Rec8 bands are indicated with red and green bars on the left. **B**. Quantification of Rec8 band intensity in (A) is shown. Rec8-enrichment to chromatin is expressed as a ratio of Rec8 to H2B levels while the soluble fraction of Rec8 is based on the ratio of Rec8 to tubulin levels. Rec8 level in *cdc20-mn* strain whole cell extracts (W) at 5 h was used to normalize the values. *P*-values were obtained by comparing signal of Rec8 in chromatin-unbound fraction (S) between 5 and 8 h. Error bars show the S.D. (n=3). **C**. Intensity of acetyl-Smc3 shown in (A) was quantified and analyzed as described in (B). **D**. Single culture of *cdc20-mn pCUP-Esp1-9myc* (KSY1009/1010) strain was synchronized and divided into two cultures; then Esp1 expression was induced by addition of 50 µM CuSO4 at 3 and 5 h. Prepared chromosome spreads were immuno-stained for Rec8 (red) and Ctf19 (green). Representative images are shown. The bar indicates 2 µm. **E**. Total Rec8 and DAPI signal intensity was quantified as shown in Fig. 1C. A minimum 30 nuclei were quantified in each representative time points. Error bars show the S.D. (n=3). **F**. Chromatin fractionation assay of *cdc20-mn pCUP-Esp1-9myc* (KSY1009/1010) cells without and with overexpression of Esp1 was carried out as shown in (A). **G**. Quantification of Rec8 levels in *cdc20-mn pCUP-ESP1-9myc* (KSY1009/1010) strain was analyzed as shown in (B). Whole cell extracts (W) at 5 h sample was used for normalization.

We next checked the status of Smc3 acetylation at K112 and K113 during meiosis by chromatin fractionation [11]. Acetylation of Smc3 occurs during DNA replication in order to facilitate the establishment of SCC. In *cdc20mn* mutants, most of the acetylated Smc3 was found in the chromatin-bound fractions at 5 h. This further confirmed that SCC formation is mediated by Smc3 acetylation in prophase-I. At 8 h, however, 51.6% of acetylated Smc3 was recovered in the unbound fraction (Fig 2A and 2C, and S1A Fig). This showed that not only Rec8, but also acetylated Smc3, a core component of the cohesion complex, is released from the chromatin in late meiotic prophase-I. Consistent with this, in *Xenopus* egg extracts cohesin release by the prophase pathway also decreased the level of acetylated SMC3 that was chromatin bound [15].

To see if the chromatin-bound fraction of Rec8 during late prophase-I remains sensitive to removal by separase, we artificially induced expression of the separase, Esp1, in prophase-I under conditions that deplete Cdc20 (*cdc20-mn*). Esp1 expression was driven by the *CUP1* promoter and copper was added at 3 and 5 h to induce Esp1 during prophase-I [22]. As above, Rec8 staining was reduced on chromatin at 8 h without Esp1 induction. Following Esp1 induction, a large number of Rec8 foci/lines disappeared, leaving only a few Rec8 foci per chromosome (Fig 2D). Indeed, the signal intensity of Rec8 dropped to 17.7±6.7% upon Esp1 induction, while at 5 h without Esp1 induction the Rec8 signal was reduced to 50.8±21.5% (Fig 2E). This demonstrated that most of Rec8 on chromatin during late prophase-I in *cdc20mn* cells remained sensitive to separase. Similar results were obtained when copper was added at 3 h. Finally, the sensitivity of Rec8 to Esp1 was confirmed by chromatin fractionation (Fig 2F and 2G, and S1B Fig). After 3 h of separase induction (8 h), the amount of full length Rec8 on chromatin was reduced to 19.5±5.9% (versus approximately 47.1±16.2% without Esp1 induction). The cleaved Rec8 products were too unstable to detect without a *ubr1* mutation which protects the product from degradation [37]. Separase-resistant Rec8 foci often co-localize with the centromere marker Ctf19 (Fig 2D). This confirms that kinetochores in late-prophase-I are able to protect Rec8 cohesin from cleavage by separase, while the arm-bound Rec8 seems to be more sensitive.

### Rec8 phosphorylation is associated with dissociation of Rec8 at late prophase-I

To characterize the phosphorylation status of Rec8 during late prophase-I, we used two phospho-specific antibodies; anti-Rec8-pS179 (PLK site) and anti-Rec8-pS521 (DDK site; kindly gifted by A. Amon, MIT) [24, 29]. Probing chromatin fractions revealed that Rec8-pS179-specific signals were very few at 5 h, but increased at 8 h in whole cell lysates from the *cdc20mn* mutant. The signals were nearly undetectable in *cdc5mn cdc20mn* mutants, illustrating dependence on Cdc5 (Fig 3A and 3B). Importantly, at 8 h, the Rec8-pS179 signal was recovered primarily in chromatin unbound fraction. This indicated that Cdc5-dependent S179 phosphorylation is associated with Rec8 release from meiotic chromatin. We were unable to detect the Rec8-pS179 signal on spreads. This may either indicate that the Rec8-pS179 signal was removed from spreads, or that the Rec8-pS179 antibody is too weak to detect the signal on spreads.

**Figure 3.**
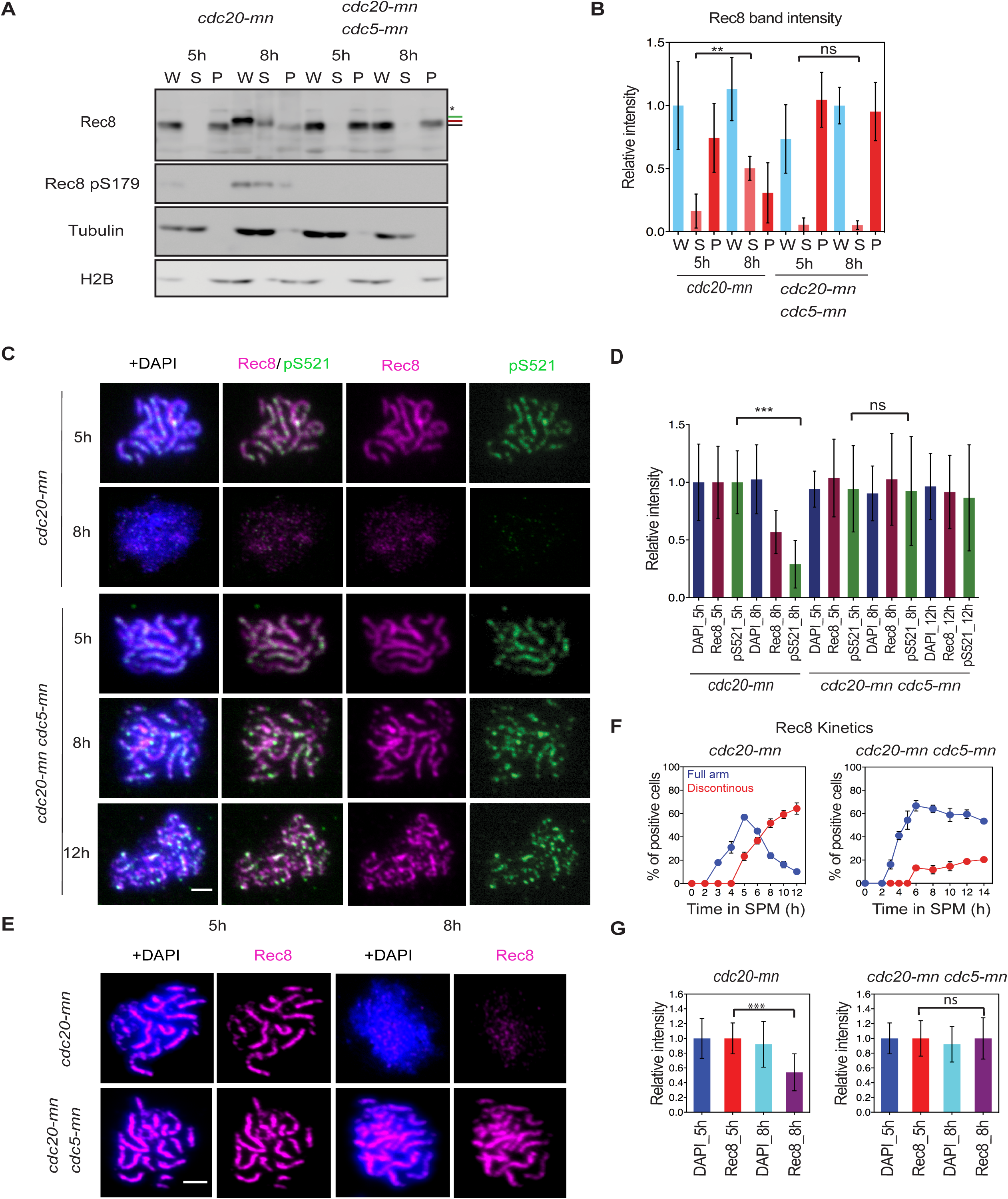
Rec8 phosphorylation is required for efficient dissociation of Rec8 at late prophase-I. **A**. Chromatin fractionation assay for *cdc20-mn* (KSY642/643) and *cdc20-mn cdc5-mn* (KSY659/660) mutant cells was carried out as described in Fig. 2A. **B**. Quantification of Rec8 band intensity in (A) is performed as shown in Fig. 2B. Error bars show the S.D. (n=3). **C**. Localization of Rec8 (red) and Rec8-pS521 (phospho-S521; green) was analyzed in *cdc20-mn* (KSY642/643) *cdc20-mn cdc5-mn* (KSY659/660) cells at 5, 8 and 12 h. **D**. Total Rec8, Rec8-pS521, and DAPI signal intensity was studied as in Fig. 1C. Error bars show the S.D. (n=3). **E**. Localization of Rec8 (red) with or without DAPI (blue) in *CDC20-mn* (KSY642/643) and *CDC20-mn CDC5-mn* (KSY659/660) mutants. Representative image is shown. The bar indicates 2 µm. **F**. Kinetics of Rec8 in (E) was classified as shown in Fig. 1B. Error bars show the S.D. (n=3). **G**. Total Rec8 and DAPI signal intensity was quantified as shown in Fig. 1C. A minimum 30 nuclei were quantified in each representative time points. Error bars show the S.D. (n=3).

We stained meiotic chromosome spreads with anti-Rec8-pS521 antibody (Fig 3C) [24], and, similar to Rec8 staining result, Rec8-pS521 shows a linear signal at 5 h in the *cdc20mn*. However, unlike Rec8, some of Rec8-pS521 (a target of DDK) foci are brighter than other foci or lines, suggesting local enhancement of S521 phosphorylation. Importantly, Rec8-pS521-specific signal on spreads was reduced at 8 h, leaving several bright foci. This loss of Rec8 signal depends on Cdc5 kinase since *cdc5mn cdc20mn* maintained high levels of Rec8-pS521 on chromosomes at the late time points (Fig 3C and 3D). Quantification revealed that Rec8-pS521-specific signal was even more strongly reduced than the global Rec8 signal (28.9±20% versus 56.8±18.6%). This is consistent with a model whereby Rec8 phosphorylation at S521, possibly by DDK, may play a role in PLK-dependent cohesin release. We were unable to detect pS521 signals on western blots efficiently (S1D Fig) and therefore could not determine whether the released Rec8 is phosphorylated at S521.

Taken together, the above results showed that Rec8-cohesin dissociates from meiotic chromosomes during late-prophase-I independent of separase activation, consistent with a previous study [31]. This suggests the presence of a cleavage-independent pathway for cohesin release that correlates with Rec8 phosphorylation. This is similar to phosphorylation-dependent cohesin release in vertebrate pro-metaphase, the so-called mitotic “prophase pathway” [14].

### Rec8 phosphorylation is required for efficient dissociation of Rec8 at late prophase-I

To examine the role of Rec8 phosphorylation in cohesin release during late prophase-I, we localized phosphorylation-deficient Rec8 mutant proteins, Rec8-17A and -29A on meiotic spreads [24, 38]. We introduced *rec8-17A* and *-29A* mutations (S2A Fig) into the *cdc20mn* background. The Rec8-17A still can be phosphorylated and shows a band shift, while Rec8-29A does not (S2B Fig). We performed the same Rec8 localization studies and Rec8 staining was characterized as linear or discontinuous as above, corresponding to the pachytene and late (early) prophase-I stages, respectively. Both Rec8-17A and -29A proteins showed linear staining patterns on meiotic chromosomes, like the wild-type Rec8 protein (Fig 4A and 4B, and S2C Fig). In *rec8-17A* mutants, the appearance of discontinuous Rec8 staining at late time points was slightly delayed as compared with the control (S2D Fig). *rec8-29A* mutants, on the other hand, showed a strong delay in the appearance of discontinuous dots of Rec8 staining, and of the subsequent dissociation of cohesin (Fig 4E and 4F). This was confirmed by intensity measurements (Fig 4G and S2E Fig). The *rec8-29A* mutant retained 77.6±25.7% of its Rec8 signal at 10 h when compared with the level at 6 h. However, because the significant delay we observed for the release of this Rec8 mutant in late prophase-I might be due to defects in earlier events, such as the processing of meiotic recombination intermediates [38], we also localized Rec8-29A mutant proteins on chromosome spreads that lacked Red1 signals, which correspond to late prophase-I. The change in Red1 staining showed a delay in late prophase-I in the *rec8-29A* mutant (Fig 4A and 4B). A delay in SC disassembly was also observed in the mutant (Fig 4C and 4D). Importantly, we found that 67.4% (n=68) of Red1-negative nuclei showed linear Rec8 staining in the *rec8-29A cdc20mn* mutant, a value that was only 5.17±0.75% (n=76) in Rec8+ *cdc20mn* cells. It is thus very likely that Rec8 phosphorylation promotes the dissociation of Rec8 cohesin from the chromosomes in late prophase-I.

**Figure 4.**
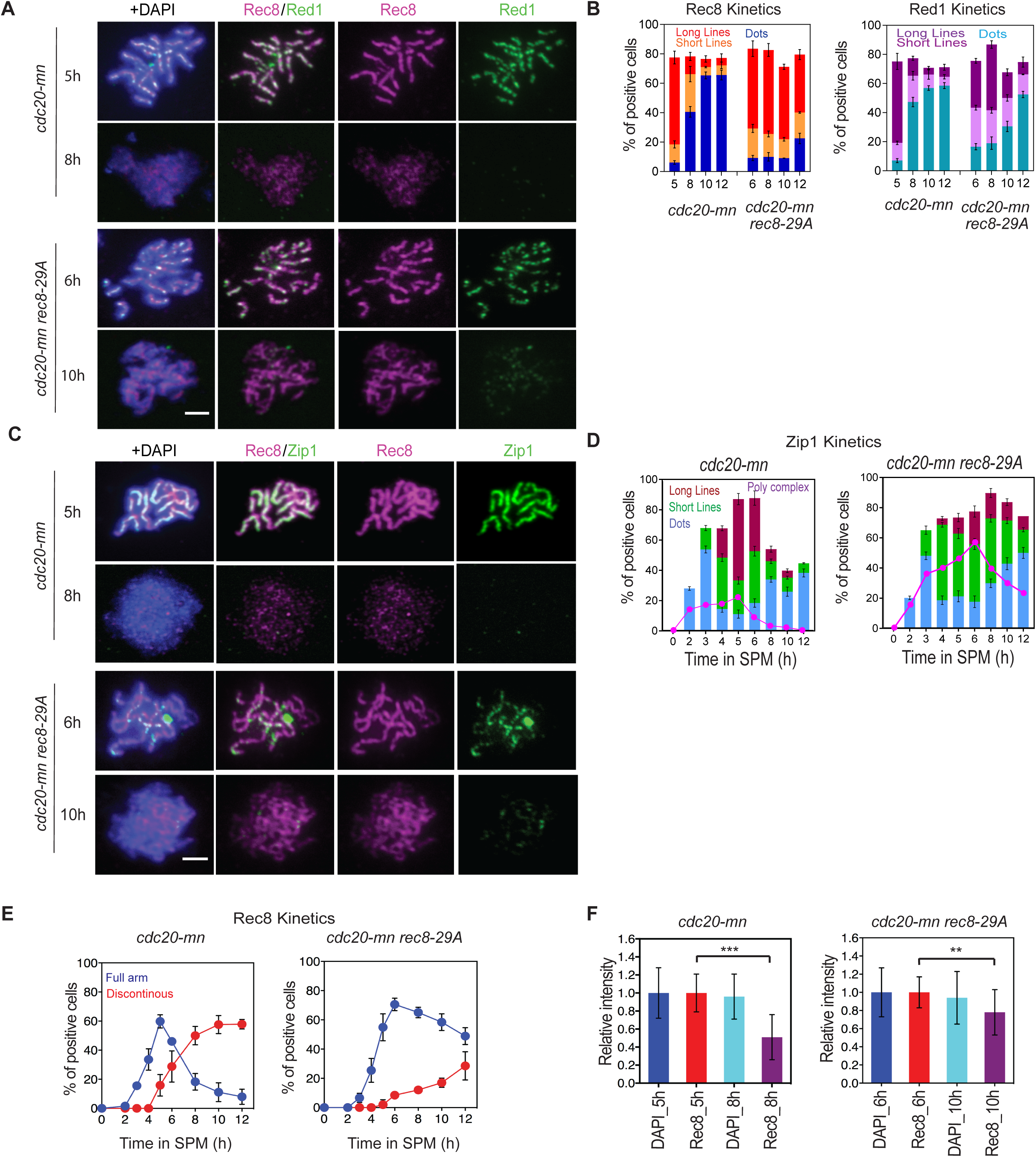
Rec8 phosphorylation is critical for cleavage-independent Rec8 dissociation from meiotic chromosomes. **A**. Localization of Rec8 (red) and Red1 (green) on chromosome spreads was analyzed for *CDC20-mn* (KSY642/643) and *CDC20-mn rec8-29A* (KSY866/867) cells. Representative image with or without DAPI (blue) dye is shown. The bar indicates 2 µm. **B**. Kinetics of Rec8 and Red1 staining classes in (A) was analyzed as in Fig. 1B. A minimum 100 cells were analyzed at each time point. Error bars show the S.D. (n=3). **C**. Localization of Rec8 (red) and Zip1 (green) spreads was studied in *CDC20-mn* (KSY642/643) and *CDC20-mn rec8-29A* (KSY866/867). **D**. Kinetics of Zip1 staining in (F) was classified shown as in Fig. 1B. Error bars show the variation from two independent experiments. **E**. Kinetics of Rec8 (left) in (E) was analyzed as in Fig. 1B. **F**. Total Rec8 and DAPI signal intensity was quantified as described in Fig. 1C. Error bars show the S.D. (n=3).

### Cdc5 is indispensable for cleavage-independent Rec8 dissociation from meiotic chromosomes

During the vertebrate mitotic “prophase pathway” for cohesin release is strongly dependent on Polo-like kinase (PLK) and other kinases [14]. Therefore, we wondered whether Cdc5, the budding yeast PLK, also regulates this release through phosphorylation of Rec8, as previously suggested [31]. We depleted Cdc5 during meiosis in the absence of Cdc20 (*cdc5mn cdc20mn)*. Indeed, Cdc5 depletion greatly reduced the appearance of discontinuous dots of Rec8 staining, and preserved Rec8 intensity at later time points, such as at 8 h (Fig 3E, 3F and 3G). This was supported by chromatin fractionation (Fig 3A and 3B). At 8 h, approximately half of Rec8 was released from chromatin in *cdc20mn* cells, and the release was not seen at 8 h in *cdc5mn cdc20mn* mutants (80±16% at 8 h relative to 5 h). In the absence of Cdc5, the mobility of Rec8 was rarely shifted up (Fig 3A). Therefore, we conclude that Cdc5/PLK is critical for cleavage-independent removal of Rec8 in late meiotic prophase-I, possibly through the phosphorylation of Rec8. Alternatively, in the absence of Cdc5, chromatin might be more highly compacted than in a normal meiosis [39], indirectly affecting the dissociation of the cohesin.

A previous study showed that ectopic expression of Cdc5 is sufficient for exit from the mid-pachytene stage in *ndt80* mutants. This is triggered by resolution of recombination intermediates into products, as well as the disassembly of SC (without entry into meiosis I) [39]. We next asked whether Cdc5 is sufficient for Rec8 chromatin dissociation by expressing Cdc5 ectopically during an *ndt80*Δ arrest. Expression of Cdc5 was induced by the addition of estradiol into the *CDC5-in ndt80* strain (Fig 5C). In concert with Zip1-disassembly, Cdc5 induction led to the formation of discontinuous Rec8 staining and thus cohesin release (Fig 5A and 5B). This process was dependent on the kinase activity of Cdc5, as kinase-dead *CDC5_kd_* (*CDC5-N209A*) mutants did not induce remodeling of the Rec8-containing structure during pachytene (Fig 5A and 5B). Our data argue that Cdc5 is both necessary and sufficient for cohesin release in mid/late pachytene.

**Figure 5.**
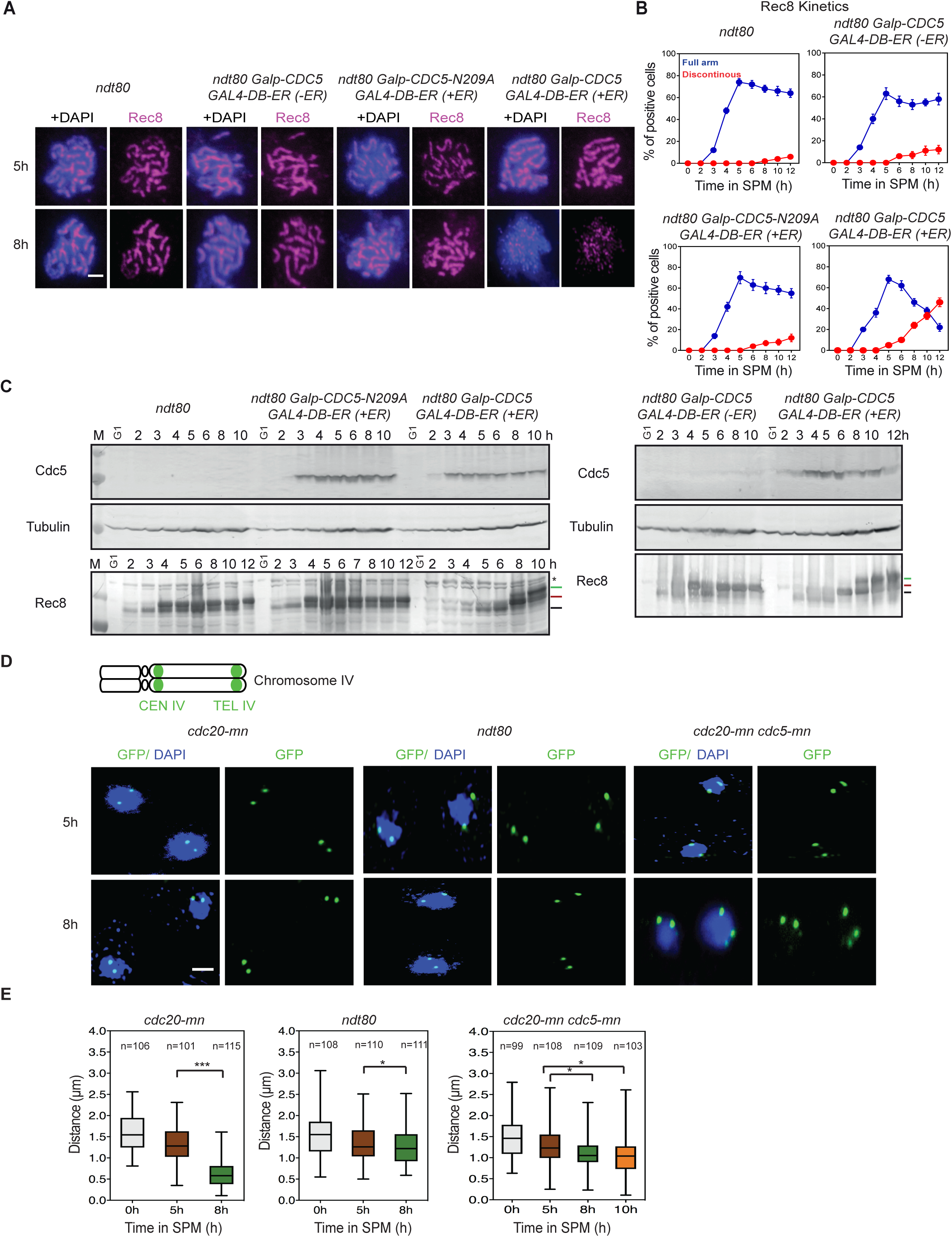
Cdc5 is sufficient for cleavage-independent Rec8 dissociation from meiotic chromosomes. **A.** Localization of Rec8 (red) with or without DAPI (blue) in *ndt80* (KSY467/468), *ndt80 GALp-CDC5 GAL4-DB-ER* without estradiol induction (-ER) (KSY887/888), *ndt80 GALp-CDC5-N209A GAL4-DB-ER* with estradiol induction (+ER) (KSY882/883) and *ndt80 GALp-CDC5 GAL4-DB-ER* (-ER) (KSY887/888) cells is shown. *CDC5* overexpression was induced by the addition of 400 nM Estradiol at 2 h. **B**. Kinetics of Rec8 classes in (A) was classified shown in Fig. 1B. A minimum 100 cells were counted at each time point. **C**. Expression profiles of Cdc5 during meiosis were verified by western blotting in *ndt80* (KSY467/468), *ndt80 Galp-CDC5 GAL4-DB-ER* without estradiol induction (-ER) (KSY887/888), *ndt80 GALp-CDC5-N209A GAL4-DB-ER* with estradiol induction (+ER) (KSY882/883) and *ndt80 GALp-CDC5 GAL4-DB-ER* (-ER) (KSY887/888) cells is shown. *CDC5* overexpression was induced by the addition of 400 nM Estradiol at 2 h. At each time point of meiosis, cells were fixed with trichloroacetic acid (TCA), and cell lysates were examined. Tubulin used was utilized as a loading control. The protein positions are indicated with the lines on the right. **D**. Representative images of *CEN4* and *TEL4* GFP foci (green) and nuclei (blue) in *cdc20-mn* (KSY991/642), *ndt80* (KSY445/467) and *cdc20-mn CDC5-mn* (KSY989/659) cells in a single focal plane of whole cell staining at each time point are shown. The bar indicates 2µm. **E**. Distances between *CEN4* and *TEL4* at each time point 0h, 5h, and 8h (*cdc20-mn;* KSY991/642 and *ndt80;* KSY445/467) as well as at 10 h in *cdc20-mn CDC5-mn* (KSY989/659) were measured and plotted as a box/whisker plot. A minimum 100 nuclei were studied in each time point.

### Rad61/Wpl1, the Wapl ortholog in yeast, regulates cohesin release during late-prophase-I

In the budding yeast, cohesin association in the mitotic G1 phase is inhibited by a Wapl ortholog, Rad61/Wpl1 [17]. The anti-cohesin activity of Rad61/Wpl1 is counteracted by Eco1-dependent acetylation of Smc3 [5, 13] and no prophase-like pathway has been reported for G2 phase in budding yeast mitosis [17]. In mammalian cells, on the other hand, a vertebrate-specific protein, sororin, counteracts Wapl activity [8]. The fact that the budding yeast does not possess a sororin ortholog prompted us to examine the role of Rad61/Wpl1 in cohesin release during meiosis. Indeed, our previous report showed that in the *rad61/wpl1* mutant, the disassembly of Rec8 is much slower than the other axis component, Red1, whose disassembly is tightly correlated with Rec8 in wild-type [40]. This suggested an uncoupling of disassembly steps for the two axis components during late prophase-I in *rad61/wpl1* mutants. Localization of Rec8 was examined in *cdc20mn* mutants lacking *RAD61/WPL1* (Fig 6A and 6B). As compared with *cdc20-mn, rad61/wpl1 cdc20mn* cells showed prolonged persistence of Rec8 lines at very late time points. Even at 14 h, 32±3% *rad61/wpl1* cells retained full linear Rec8 staining (Fig 6A and 6C). Indeed, the signal intensity of Rec8 was unchanged between 5 and 8 h in the absence of Rad61/Wpl1 (Fig 6B). This suggests a key role of Rad61/Wpl1 in cohesin release in the G2 phase of yeast meiosis. Moreover, decrease of phosphorylated S521 of Rec8 signals during late prophase-I is largely dependent on *RAD61/WPL1* (S1C Fig), suggesting that Rad61/Wpl1 is critical for the release of phosphorylated Rec8 from the chromosomes.

**Figure 6.**
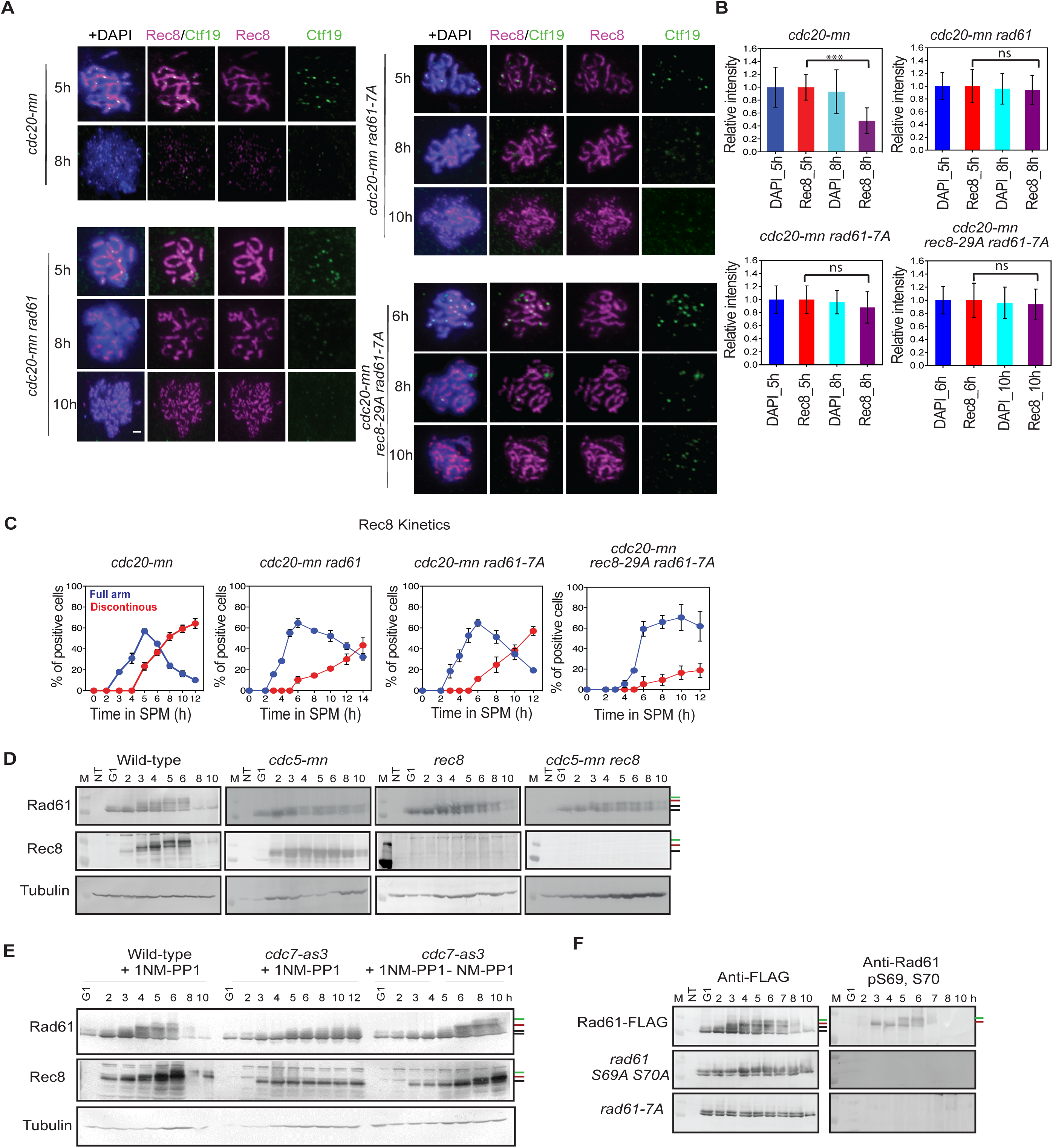
Rad61/Wpl1 plays a role in cohesin-release in late prophase-I. **A.** Localization of Rec8 (red) and Ctf19 (green) in *cdc20-mn* (KSY642/643), *cdc20-mn rad61* (KSY637/638), *cdc20-mn rad61-7A* (KSY653/654) and *cdc20-mn rec8-29A rad61-7A* (KSY1043/1044) cells. Representative image for each strain at various time points are shown. **B**. Measurement of total Rec8 and DAPI signal intensity in (a) was analyzed in Fig. 1C. Error bars show the S.D. (n=3). **C**. Kinetics of Rec8-classes was studied as described in Fig. 1B. A minimum 100 cells were analyzed per time point. Error bars show the S.D. (n=3). **D**. Expression profiles of Rec8 and Rad61-Flag during meiosis were verified using *RAD61-FLAG* (KSY440/441), *cdc5-mn RAD61-FLAG* (KSY434/435), *rec8 RAD61-FLAG* (KSY627/628) and *rec8 cdc5-mn RAD61-FLAG* (KSY1092/1093) cells by western blotting. The protein positions are indicated with the lines on the right. The positions of Rec8 and Rad61-Flag are shown by bars. Black bars indicate non-phosphorylated Rad61 or Rec8. Red and green bars are DDK-dependent and PLK/Cdc5-dependent phosphorylated Rad61 or Rec8, respectively. Bands shits of Rec8 and Rad61 were analyzed in presence and absence of 1NM-PP1 in *RAD61-FLAG* (KSY440/441) and *cdc7-as3 RAD61-FLAG* (KSY978/979) strains at the indicated time points by western blotting as shown in (D). 1NM-PP1 was added at 0 h in three cases and washed out at 5 in the right panel. **E**. The western blotting analysis was carried out for Rad61-Flag (Right) and Rad61-pS69, 70 (left) in *RAD61-FLAG* (KSY440/441), *rad61-S69A, S70A-FLAG* (KSY754/757) and *rad61-7A-FLAG* (KSY753/755) strains as shown in (D).

We investigated the expression of Rad61-Flag during meiosis by western blot, and found that Rad61 exhibits multiple bands upshifted during meiosis (Fig 6D). Similar to Rec8, Rad61 expression decreases after 8 h. In addition to the two bands observed during pre-sporulation at 0 h, at least two major meiosis-specific forms of Rad61 were observed; one that started to appear at 3 h, and a second that appeared at 5 h. The slowly migrating forms of Rad61 disappeared at 8 h and were far less abundant relative to early time points. The appearance of two meiotic-specific forms of Rad61 protein, as well as its disappearance resembles Rec8, which also showed two major phosphorylated species in addition to the unmodified one [24, 25]. It seemed likely that Rad61 band shifts were due to phosphorylation, and since Rec8 phosphorylation is catalyzed by three kinases, DDK, PLK, and CK1 [24, 25], we checked the effects of these kinases on Rad61 modification. When the kinase activity of analog-sensitive Cdc7 (Cdc7-as3) was suppressed by its inhibitor, PP1, band shifts of both Rec8 and Rad61 were greatly diminished in meiosis (Fig 6E). After washing out PP1, the band shifts reappeared. This indicated that band shifts of Rad61 are dependent on Cdc7 (DDK) kinase activity.

We also checked the depletion of Cdc5 and found that the upper bands of both Rad61 and Rec8 at late time points (5 and 6 h) were nearly abolished in the *cdc5-mn* cells (Fig 6D). These results showed that, like Rec8, the Rad61-band shift requires both DDK and PLK activities. Rad61 phosphorylation is independent of meiotic recombination and DSB formation, since *spo11-Y135F* mutants displayed normal Rad61 and Rec8 band shifts (S3A Fig). On the other hand, Rec8 is essential for the PLK(Cdc5)-dependent secondary band shift of Rad61, although not the DDK-dependent one (Fig 6D). As a control, we also found band shifts of Rad61 in *rec8 cdc5-mn* cells similar to those in *rec8* cells (Fig 6D). This is consistent with the fact that Rec8 directly binds to Cdc5 kinase [25].

Based on the sequence information of Rad61 [41], we mapped putative DDK sites in the N-terminal non-conserved region of Rad61, outside of the conserved WAPL domain (S4A Fig). These sites were as follows: T13, S25, S69, S70, T95, S96, and S97. Various substitution combinations were generated for these putative sites: *rad61-T13A, S25A-FLAG, rad61-S69A, S70A-FLAG, rad61-T95A, S96A, S97A-FLAG, rad61-T13A, S25A, T95A, S69A, S70A, S96A, and S97A-FLAG* (hereafter, *rad61-7A*) and we found that meiosis-specific band shifts of Rad61 were compromised in the *rad61-S69A, S70A-FLAG* and *rad61-7A* mutants but not in the *rad61-T13A, S25A-FLAG* and *rad61-T95A, S96A, S97A-FLAG* strains (Fig 6F and S4B Fig). We raised an antibody against a Rad61 peptide containing phospho-S69 and phospho-S70 sites and used it to detect phosphorylation-specific bands of Rad61. Western blotting using Rad61 phospho-specific antibody clearly revealed two meiosis-specific bands, which were absent in mitosis (Fig 6F, right panels).

The *rad61-7A* mutant exhibited spore viability comparable to wild-type cells (S4C and S3D Fig), and entry into meiosis I was only delayed one hour (S3B Fig), suggesting that Rad61 phosphorylation plays a minor role in early prophase-I. We then investigated the effect of the *rad61-7A* mutation on Rec8 dissociation at late prophase-I in the absence of Cdc20. Compared to the *cdc20mn* mutant, the *rad61-7A cdc20mn* mutant showed a delayed disappearance of the linear staining and an appearance of discontinuous Rec8 dots at later time points (Fig 6A and 6C). Again, linear Rec8 expression was frequently detected in Red1-negative nuclei of *rad61-7A cdc20mn* cells (Fig 6A). The defective Rec8 release in *rad61-7A* mutants was also confirmed by Rec8-intensity measurements (Fig 6B).

This *rad61-7A* defect resembles the *rec8-29A* mutant, although it is less pronounced than the *rad61* null mutant phenotype. The *rad61-7A rec8-29A* double mutant showed an even more delayed disappearance of Rec8 lines than the two single mutants (Fig 6A and 6C), indicating that both Rec8 and Rad61 phosphorylation contribute to Rec8 release in late prophase-I. The *rad61-7A* and the *rec8-29A* single mutant show 94% and 72.7% spore viability, respectively, while spore viability in the *rad61-7A rec8-29A* double mutant was reduced to 64.1% (S4C and S4D Fig). This suggests that the phosphorylation-triggered release is physiologically relevant for meiotic progression.

Finally, we checked a chromosome segregation defect in the mutant deficient in cohesin release by using *CENV-GFP* [42]. The *rad61-7A, rec8-29A* and *rad61-7A rec8-29A* mutants are proficient in sister chromatid cohesion during prophase-I (S4E left, Fig). Although the *rad61-7A* and *rec8-29A* single mutants showed little defect in disjunction of homologous chromosomes at meiosis I, the *rad61-7A rec8-29A* double mutant showed slight, but significant increase of mis-segregation of the chromosomes (*P*=0.013; S4E right, Fig). These results support the notion that Rad61 and Rec8 phosphorylation play a redundant role for the segregation of homologous chromosome by regulating the phosphorylation status of cohesin components such as Rec8 and Rad61/Wpl1.

### PLK promotes chromosome compaction in late prophase-I

To observe the consequences of Rec8-cohesin release in late prophase-I, we measured chromosome compaction using two fluorescently marked chromosome loci on chromosome IV in three strains, *ndt80, cdc20mn*, and *cdc5mn cdc20mn* (Fig 5D). At 0 h, the distance between the two loci was 1.55±0.42 μm in the *cdc20mn* mutant, and this was reduced to 1.3±0.4 μm at 5 h (Fig 5E) with no further decrease at 8 h in the *ndt80* mutant (1.3±0.5 [5 h] and 1.2±0.4 μm [8 h]). This is consistent with a compaction that occurs in pachytene chromosomes [40]. The *cdc20mn* mutant showed an additional decrease of the distance to 0.6±0.3 μm at 8 h, or compaction to 37% of initial length. This argues for a specific chromosome compaction event of ∼3-fold in prophase-I after pachytene stage. Importantly, this drastic chromosome compaction completely depends on Cdc5 PLK. The *cdc5mn cdc20mn* cells showed only mild compaction of chromosomes both at 5 and 8 h (Fig 5D and 5E).

## Discussion

Together with a previous study [31], our results suggest the existence of a phosphorylation-controlled step during late prophase-I/pro metaphase-I that leads to the partial release of cohesin in meiotic yeast cells. This occurs in addition to the previously identified two steps of cohesin release at metaphase/anaphase-I and -II (Fig. 7, top) [19, 37]. Prior to the final cleavage-dependent removal of cohesin, we show a cleavage-independent removal, which releases the meiotic kleisin subunit, Rec8, intact, at late prophase-I. This is a meiotic “prophase-like pathway” as it is analogous to the “prophase” pathway in mitotic G2-phase and pro-metaphase of vertebrate cells [14]. Interestingly, mitotic cells in budding yeast seem to lack the prophase pathway [17]. This is consistent with the fact that budding yeast does not possess a sororin ortholog, the key regulator of cleavage-independent removal of cohesin during the late G2 phase in vertebrates [8]. Whereas vertebrate cells inactivate the Wapl inhibitor, sororin, in mitotic prophase, meiotic yeast cells appear to regulate the activity of Wapl, Rad61/Wpl1, positively by meiosis-specific phosphorylation, and also to control Rec8’s affinity to Smc3 negatively by meiosis-specific phosphorylation of Rec8.

**Figure 7.**
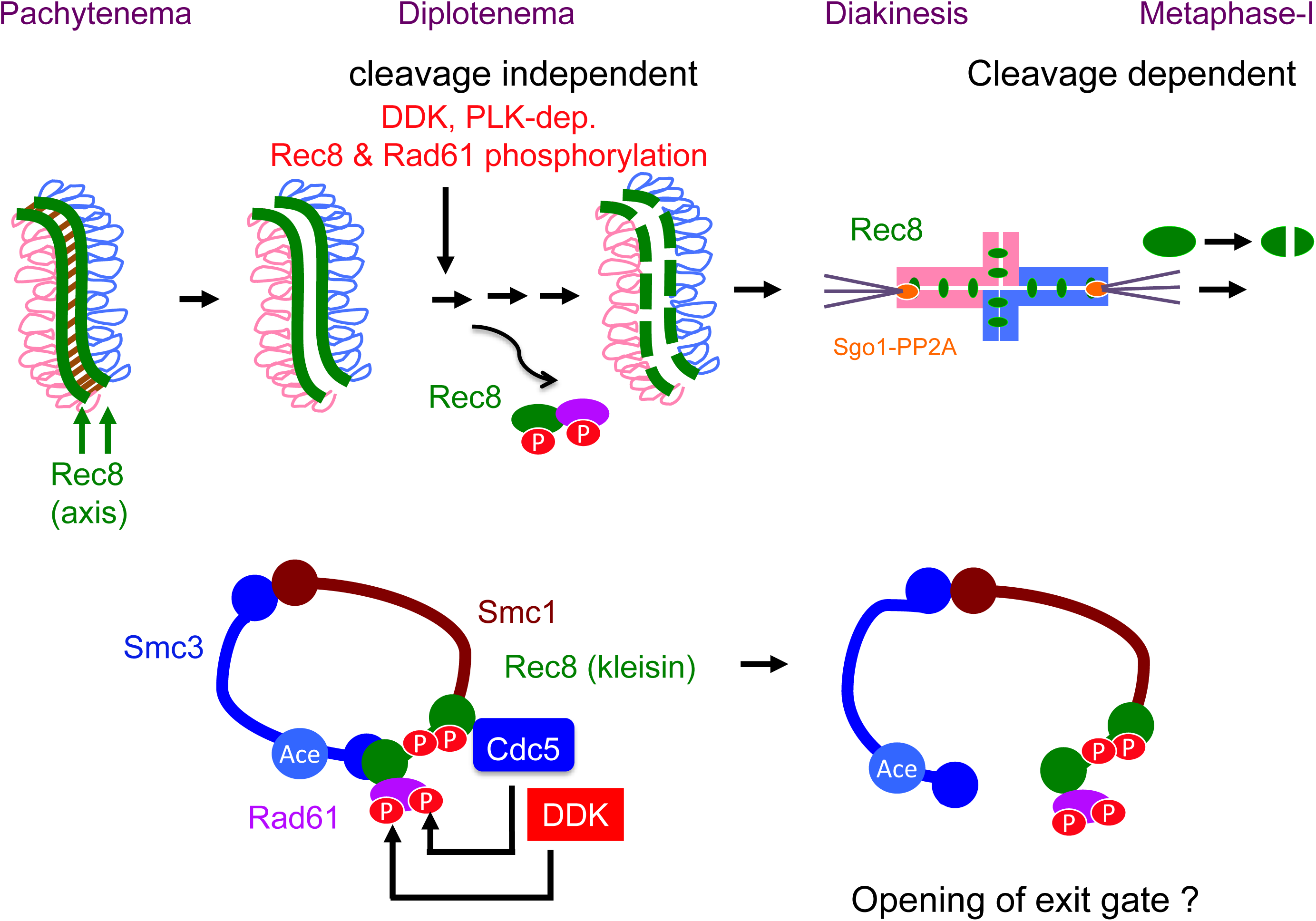
A model for meiotic prophase-like pathway at late prophase-I. Top, three step removal of cohesin during meiosis of the budding yeast. Bottom, possible model of cohesin release during late prophase-I by phosphorylation of Rec8 and Rad61, which may trigger the opening of the exit gate between Smc3 head and Rec8.

### Meiotic prophase-like pathway shares similar mechanism with the vertebrate prophase pathway for cohesin release

Similar to the vertebrate prophase pathway [14], the meiotic prophase pathway for cohesin release in budding yeast is independent of cohesin cleavage. Rec8 was released from meiotic chromosomes in the absence of separase activity in Cdc20-depleted cells, as well as in cleavage-resistant *Rec8-N* cells. Moreover, we were able to recover the full-length Rec8 protein, which was stably bound to the chromatin during mid-pachytene, in chromatin-soluble factions during late prophase-I (Fig 2A). These results show that a mechanism that releases Rec8-cohesin independent of kleisin cleavage exists.

Like the mammalian prophase pathway, the meiotic prophase pathway in yeast requires WAPL (Rad61/Wpl1) and PLK (Cdc5). During the mitotic G1 phase in yeast, Rad61/Wpl1 is known to promote the dissociation of mitotic cohesin [17]. During the mammalian mitotic prophase and the yeast G1 phase, the Wapl works together with Pds5 to mediate the opening of the exit gate between Scc1-Smc3 [11]. Judged by the role of Rad61/Wpl1, we propose that the meiotic prophase pathway is mechanistically similar to the mammalian prophase pathway and the G1 pathway in yeast. The release of meiotic cohesin in late prophase-I may occur through the opening at the interface between Rec8 and Smc3 (Fig. 7, bottom). It might be possible to confirm this by expressing a Smc3-Rec8 fusion protein that locks the interface between the meiotic kleisin, Rec8, and Smc3.

### Yeast meiotic prophase-like pathway is regulated in a different manner from the vertebrate prophase pathway

In vertebrate cells, sororin inactivation is essential for the cleavage-independent release of cohesin. Sororin is likely bound to the PDS5A/B, to which Wapl also binds. Sororin binding sterically hinders the binding of PDS5A/B to Wapl, and as a result, Wapl is unable to open the exit gate [1]. Phosphorylated sororin dissociates from PDS5A/B, allowing for binding between Wapl and PDS5A/B, which leads to opening of the gate. Given that budding yeast lacks sororin, the anti-cohesin activity of Rad61/Wpl1 in yeast is counteracted by Eco1-mediated acetylation of Smc3, which is sufficient to antagonize Rad61 activity in G2 phase [5, 13]. We found that Smc3 acetylation is maintained during prophase-I (G2 phase) of meiosis (Fig 2A). Thus, rather than inactivating a negative regulator for Wapl, yeast meiotic cells seem to display a novel mechanism for cohesin release through enhancement of Rad61/Wpl1 activity. This enhancement correlates with the phosphorylation of the Rad61, which may either enhance Wapl activity directly and/or increase the interaction between Rec8 and Rad61.

In addition to previously identified Cdc5/PLK [31], we identified two critical regulators of the meiosis prophase pathway, Rad61 and DDK. All three regulators are expressed during mitosis and meiosis. Nevertheless, our results show a meiosis-specific regulation of cohesin removal. Since, during meiosis, Scc1 is replaced by the meiosis-specific kleisin Rec8, we propose that Rec8 is an essential component for the meiosis-specific prophase pathway. In mitotic cells, PLK-dependent phosphorylation of Scc1 promotes its cleavage, rather than the release. Thus, the meiotic specificity of the prophase pathway is conferred by replacing Scc1 by Rec8.

In addition, Rad61 is phosphorylated only in meiotic prophase-I by the two mitotic kinases, DDK and PLK. Rec8 is known to interact directly with Cdc5/PLK [25]. We also show that the meiosis-specific Cdc5-dependent phosphorylation of Rad61 requires Rec8. Therefore, Rec8 has dual functions in cohesin release during meiosis. Rec8 exhibits an intrinsic property that allows it to respond to the anti-cohesin activity of Wapl, as well as the ability to enhance Wapl activity by promoting its phosphorylation. The exact mechanism that induces the meiosis-specific, DDK-dependent phosphorylation of Rad61 in early prophase-I is unknown. We know that Rec8 at least does not play an essential role in this phosphorylation, as *rec8* mutants were able to initiate meiosis-specific DDK-dependent phosphorylation of Rad61. We propose that phosphorylation of Rad61 may augment anti-cohesin activities, and consequently, the gate-opening activity of the protein (Fig. 7). In addition, we also propose that Rec8 phosphorylation may loosen the binding of Rec8 to Smc3 and Rad61 to unlock the Rec8-Smc3 gate.

### The meiotic prophase pathway is conserved in higher eukaryotes

The cleavage-independent pathway of cohesin release during meiosis is conserved in higher eukaryotes such as nematodes, plants, and mammals. In these organisms, differential distributions and/or reduced signals of cohesin on chromosomes or chromosome arms were observed in late prophase-I, as in diakinesis. In nematodes, cohesin on short arms, but not on long arms, is likely to be removed, in a manner dependent on aurora kinase, *air-2* [43]. Interestingly, in nematodes, Wapl (*wapl-1)*, controls the dynamics of kleisin COH3/4-containing cohesin, but not of cohesin associated with Rec8 [44]. Immuno-staining showed that in *Arabidopsis thaliana*, most Rec8 molecules are released from meiotic chromosomes during the diplotene stage, and this is mediated by Wapl [45]. Like yeast, *C. elegans* and *A. thaliana* lack a clear sororin ortholog, indicating that the removal of cohesin during meiosis is sororin-independent.

Rec8 is conserved from yeasts to mammals. Thus, the meiotic prophase pathway might be also conserved in mammals. On the other hand, in mammals, in addition to REC8, the other meiosis-specific kleisin, RAD21L, and also meiosis-specific SMC1β and STAG3 are expressed [46]. Therefore, the control of cohesin release during mammalian meiosis may be more complicated. In mouse spermatocytes, the RAD21L kleisin, but not REC8, is predominantly removed during the diplotene stage, in a manner partially dependent on PLK [18]. Recently, a novel regulatory circuit for cohesin removal was described during spermatogenesis, where NEK1 kinase-dependent “de”phosphorylation of WAPL promotes its retention on chromosomes and consequently the release of cohesin [47]. It was observed that during meiosis in mouse spermatocytes, phosphorylation of Wapl inhibits its activity. This is in sharp contrast to the role of Rad61 phosphorylation in the budding yeast, but may again reflect the absence of sororin in yeast.

### Local regulation of protection and promotion of cohesin removal along the chromosomes

Results presented in this work showed that approximately 50-60% of chromosome-bound Rec8 at the pachytene stage can be dissociated from chromosomes during late prophase-I. On the other hand, 40-50% of Rec8 remains stably bound to chromosomes during late prophase-I, suggesting that these Rec8 molecules are either protected against or are not activated for the meiotic prophase-like pathway. Most of the chromatin-bound Rec8 at late prophase-I is still sensitive to artificially expressed separase while, as expected, Rec8 at kinetochores is resistant to it. At the onset of anaphase-I, kinetochore-bound Rec8 is protected by a molecule called Shugoshin (Sgo1) [25, 27], which is bound to kinetochores during late prophase-I. Indeed, artificial expression of separase in pachytene-arrested cells; e.g. *ndt80* mutant, completely removes Rec8 even at kinetochores [22]. Thus, the full protection of kinetochore-bound Rec8 is established only after the exit from mid-pachytene. In the mammalian prophase pathway, kinetochore-bound cohesin is protected by Shugoshin/PP2A which dephosphorylates subunits like sororin. A similar protection mechanism seen in the mammalian mitotic prophase pathway may also operate on cohesins that are bound to meiotic chromosome arms and to the kinetochores. However, protection of arm cohesin, which is sensitive to separase, must be functionally distinct from kinetochore-cohesin, which is not.

The meiotic prophase pathway requires DDK- and PLK-dependent meiosis-specific phosphorylation of Rec8 and Rad61. Indeed, we showed that the Rec8 released from chromosomes is more phosphorylated than the complement that remains tightly bound to chromosomes. One plausible mechanism for the observed protection against the prophase pathway is local activation of dephosphorylation of cohesin, as seen at kinetochores, where Shugoshin recruits the phosphatase PP2A. This is similar to the role that Sgo1 plays in protection of centromeric cohesin at the onset of anaphase-I in vertebrate meiosis.

Alternatively, local activation by phosphorylation of Rec8 and Rad61 may be a mechanism that promotes cohesin release in distinct chromosomal regions. For instance, local removal of cohesion at the site of chiasmata [48] may be a necessary step in the formation of normal diplotene bivalents.

### Is Rec8 phosphorylation indeed required for the cleavage by separase?

Previous reports strongly suggested that phosphorylation of Rec8 by DDK, PLK, and CK1 is essential for cleavage by separase [24, 25]. However, this was not directly tested by an *in vitro* cleavage assay. Our results presented here suggest an additional role for Rec8 phosphorylation by DDK and PLK: the dissociation of Rec8-cohesin at late prophase-I. However, we and others [22] also showed that chromosome-bound Rec8 with reduced phosphorylation can be a substrate for separase-mediated cleavage *in vivo*, since ectopic expression of separase in late prophase-I was sufficient for the removal of Rec8-cohesin from chromosome arms, but not from centromeres (Fig 2D). Thus, it is possible that phosphorylation of Rec8 plays a major role in cleavage-independent dissociation of cohesin, in addition to triggering it for cleavage by separase.

### Functions of the meiotic prophase pathway

Given that cohesin at the chromosome arms is important for chromosome segregation in MI, one may suggest that the meiotic-prophase pathway is dangerous in meiotic cells, and thus, it is important to ask why meiosis has retained this dangerous pathway. It is known that during late prophase-I, which corresponds to diplotene and diakinesis in other organisms, drastic changes in chromosome morphology occur [49]. This includes strong compaction while chiasmata emerge, to prepare for chromosome segregation. In meiosis I, the chiasmata are essential for chromosome segregation. Indeed, loss of cohesion around chiasmata sites has been observed in various organisms [48]. Similarly, in worms, Wapl-dependent cohesin removal promotes recombination-mediated change of meiotic chromosome structure [44].

We propose that one function of cohesin release in late prophase-I is to promote chiasma formation. Since the individualization of chromosomes may be important for development of chiasmata. In addition, in late prophase-I, chromosomes show drastic compaction. Even in the budding yeast, late meiotic chromosomes are compacted by approximately 3 fold as compared with their sizes in meiotic G1 (this study) [31]. Concomitant with this drastic compaction, condensin has been shown to bind to meiotic chromosomes following the pachytene stage [31, 50]. The binding of condensin not only promotes condensation, but also facilitates the release of cohesin [31], and may promote individualization of chromatids. In this work we show that Cdc5 depletion causes both a failure in the meiotic prophase pathway and a defect in chromosome compaction. This is consistent with a role of cohesin removal in compaction, although other interpretations are possible. What the basic purpose of cohesin removal in prophase is either to make space for condensin, to allow chiasma morphogenesis, or to allow compaction. Which of these, or yet another function, remains to be answered in future studies.

## Materials and Methods

### Strains and strain construction

All strains described here are derivatives of SK1 diploid strains, MSY832/833 (*MAT*α*/MAT****a***, *ho::LYS2/”, lys2/”, ura3/”, leu2::hisG/”, trp1::hisG/”*). Strain genotypes are given in S1 Table. *CEN4-GFP/TEL4-GFP* and Esp1-overexpression strains were provided by Dr. Doug Koshland and Dr. Keun P. Kim, respectively.

### Antisera and antibodies

Anti-Zip1, anti-Red1, and anti-Rec8 antisera for cytology and western blotting have been described previously [51, 52]. Secondary antibodies conjugated with Alxea488 and Alexa594 dyes (Molecular Probes, Life Technologies, UK) were used for the detection of the primary antibodies. Anti-Rec8-pS179 (PLK site) and anti-Rec8-pS521 (DDK site) were generous gifts from Dr. Angelika Amon (MIT). Anti-acetyl-Smc3 was a gift by Dr. Katsu Shirahige (U. of Tokyo). Anti-Rad61-PS69-pS70 antibody was raised in rabbit using a Rad61 peptide containing pS69 and pS70 by a company (MBL Co. Ltd).

### Cytology

Immunostaining of chromosome spreads was performed as described previously[53, 54]. Stained samples were observed using an epi-fluorescence microscope (BX51; Olympus, Japan) with a 100X objective (NA1.3). Images were captured by CCD camera (CoolSNAP; Roper, USA), and afterwards processed using IP lab and/or iVision (Sillicon, USA), and Photoshop (Adobe, USA) software tools.

### SIM imaging

The structured illumination microscopy was carried out using super resolution-structured illumination (SR-SIM) microscope (Elyra S.1 [Zeiss], Plan-Apochromat 63x/1.4 NA objective lens, EM-CCD camera [iXon 885; Andor Technology], and ZEN Blue 2010D software [Zeiss]) at Friedrich Miescher Institute for Biomedical Research, Switzerland. Image processing was performed with Zen software (Zeiss, Germany), NIH image J and Photoshop.

### Fluorescence intensity measurement

Mean fluorescence of the whole nucleus was quantified with Image J. Quantification was performed using unprocessed raw images and identical exposure time setting in DeltaVision system (Applied Precision, USA). The area of a nuclear spread was defined as an oval, and the mean fluorescence intensity was measured within this area.

### Chromatin fractionation

Chromatin fractionation was performed as described previously[55]. The cells were digested with Zymolyase 100T (Nakarai Co. Ltd) and the spheroplasts were pelleted. The pellets were resuspended in five volumes of hypotonic buffer (HB; 100 mM MES-NaOH, pH 6.4, 1 mM EDTA, 0.5 mM MgCl_2_) supplemented with a protease inhibitor cocktail (Sigma, USA). After 5 min, 120 µl of whole cell extract (WCE) were layered onto 120 μl of 20% (W/V) sucrose in HB and centrifuged for 10 min at 16,000 *g*. The supernatants were saved and the pellets were resuspended in 120 μl EBX buffer (50 mM HEPS-NaOH, pH 7.4, 100 mM KCl, 1 mM EDTA, 2.5 mM MgCl_2_, 0.05% Triton X100) and centrifuged for 10 min at 16,000 *g*. The pellets were again collected and resuspended in EBX buffer with 5 units/ml DNase I and 1 mM MgCl_2_ for 5 min. The supernatants were saved for further analysis.

### Cohesion and pairing assays

Sister chromatid cohesion and chromosome segregation during meiosis I was analysed using yeast cells heterozygous for LacI-GFP spots at *CEN5* locus [42]. Following fluorescence microscope imaging, the number of chromosomal locus-marked GFP foci in a single cell was counted manually. For sister chromatid cohesion, cells with single DAPI body at 5 h were examined. For the observations of chromosome segregation in meiosis I, cells with two DAPI bodies were selected at 6, 7 and 8 h, and the number of GFP focus in each DAPI body was counted.

### Compaction assay

For distance measurements on probed SCs at 0, 5 and 8 h, chromosome spreads were prepared as described above and stained with both anti-Rec8 and anti-GFP antibodies. The distance between two GFP foci on chromosome IV was measured by Velocity™ program (Applied Precision, USA) or IPLab (Sillicon, USA).

### Yeast culture

Yeast cell culture and time-course analyses of the events during meiosis and the cell cycle progression were performed as described previously[54].

### Statistics

Means ± S.D values are shown. Datasets were compared using the Mann-Whitney U-test. *χ*^□^-test was used for proportion. Multiple test correction was done with Bonferroni’s correction. *, **, and *** show P-values of <0.05, <0.01 and <0.001, respectively. The results of all statistical tests are shown in Supplemental Table 2.

## Supporting information

## Supplemental Information

**S1 Fig. Rec8 dissociates from meiotic chromosomes at late prophase-I. A:** Chromatin fractionation assay was carried out using *CDC20-mn* (KSY642/643) and *ndt80* (KSY467/468) mutant cells. Western blotting was performed for whole cell extracts (W), soluble fractions (S) and chromatin-bound fraction (P). Rec8 (top) and acetyl-Smc3 (second) were probed together with tubulin (third) and Histone 2B (H2B; bottom) as controls for soluble and chromatin-bound proteins, respectively. **B:** Chromatin fractionation assay of *CDC20-mn pCUP-Esp1-9myc* (KSY1009/1010) cells without and with overexpression of Esp1 was carried out as shown in (A). **C:** Localization of Rec8 (red) and Rec8-pS521 (phospho-S521; green) was analyzed in *cdc20-mn rad61* (KSY637/638) cells at 5 and 8 h. Total Rec8, Rec8-pS521, and DAPI signal intensity was studied as in Fig. 1C and shown in bottom. Error bars show the S.D. (n=3). **D:** Western blotting of Rec8 pS521 in *CDC20-mn* (KSY642/643) was done with tubulin as a control.

**S2 Fig. Rec8 phosphorylation and phosphorylation-defective mutants.** A: **A.** Schematic drawing of Rec8-17A and Rec8-29A mutant proteins. Mutated amino acid residues are shown in red. **B:** Western blotting analysis of Rec8 and tubulin was carried out using *CDC20-mn* (KSY642/643), *CDC20-mn rec8-29A* (KSY866/867) and *CDC20-mn rec8-17A* (KSY812/813) cells strain as described (A). Phosphorylated species of Rec8 and tubulin. Representative images are shown. **C:** Localization of Rec8 (red) on chromosome spreads was analyzed for *CDC20-mn* (KSY642/643) and *CDC20-mn rec8-17A* (KSY812/813) cells. Representative image with or without DAPI (blue) dye is shown. The bar indicates 2µm**. D:** Kinetics of Rec8 staining classes in (C) was analyzed as in Fig 1B. A minimum 100 cells were analyzed at each time point. **E:** Quantified total Rec8 and DAPI signal intensity was measured. A minimum 30 nuclei were quantified in each representative time points. Error bars show the S.D. (n=3).

**S3 Fig. Meiosis-specific Rad61 phosphorylation in phosphorylation-defective *rad61* mutants. A:** The western blotting analysis was carried out for Rad61-Flag in *RAD61-FLAG* (KSY440/441), *rad61-S69AS70A-FLAG* (KSY754/757) and *rad61-7A-FLAG* (KSY753/755) strains. **B:** Bands shits of Rad61 in *ndt80 RAD61-FLAG* (KSY467/468) and *spo11-Y135F RAD61-FLAG* (KSY474/475) cells were analyzed as shown in (A).

**S4 Fig. Meiotic phenotypes of phosphorylation-defective *rad61* and *rec8* mutants. A:** Schematic drawing of Rad61 with putative DDK-dependent (red) and PLK-dependent phosphorylation sites (green). Conserved “WAPL” domain is shown in a box. **B:** Kinetics of the entry into meiosis I in wild-type (MSY832/833) and *rad61-7A* (KSY753/755) cells was analyzed by DAPI counting. A cell with 2, 3, and 4 DAPI bodies was counted. At each time point, more than 100 cells were examined. **C:** Distribution of viable spores per tetrad in various strains was measured and shown. Spores were incubated after dissection at 30ºC for 3 days. Each bar indicates the percentage of classes with 4, 3, 2, 1 and 0 viable spores per tetrad. Spore viability and the total number of dissected tetrads (parentheses) are also shown. Wild type (MSY832/833), *rad61-7A* (KSY753/755), *rec8-29A* (KSY814/815), *rec8-29A rad61-7A* (KSY982/983) cells. **D:** Percentage of viable spores in various strains was shown in graph. Wild type (MSY832/833), *rad61-7A* (KSY753/755), *rec8-29A* (KSY814/815), *rec8-29A rad61-7A* (KSY982/983) cells. **E:** Sister chromatid cohesion and segregation of homologous chromosome were analyzed. A cell heterozygous for *CEN4-GFP* locus was used. At least more than 50 cells with single and two DAPI bodies in a cell were examined for the number of *CEN4*-GFP spot at 4, 5, and 6 h. For sister chromatid cohesion assay (left graph), the number of a cell containing single DAPI body with either 1 or 2 GFP spots was counted. For segregation assay of homologous chromosomes at meiosis I (right graph), a cell containing two DAPI bodies was examined for either both two DAPI bodies contained 1 GFP spot or one of two DAPI bodies contained 1 or 2 spots. Wild type (MSY833/KSY216), *rad61-7A* (KSY653/1089), *rec8-29A* (KSY814/1086), *rec8-29A rad61-7A* (KSY982/1091) cells.

**S1 Table. Strain list.** The strain used in this study and its genotype.

**S2 Table. Numerical and statistical data.** Numerical data underlying graphs and summary statistics.

## Acknowledgements

We are grateful to Drs. A. Marston, K. Matsuzaki, and H. B. D. P. Rao as well as the members of the Shinohara lab for helpful discussions. We thank Drs. A. Seeber and L. Gelman for help with SIM and Ms. A. Murakami and H. Wakabayashi for excellent technical assistance. We are grateful to Drs. A. Amon, K. Kim, D. Koshland, K. Shirahige, K. Nasmyth, and W. Zachariae for materials used in this study.

## Author contributions

**Conceptualization:** Kiran Challa, Franz Klein, Akira Shinohara

**Formal analysis:** Kiran Challa, Ghanim Fajish V

**Funding acquisitions:** Akira Shinohara

**Investigation:** Kiran Challa, Ghanim Fajish V, Miki Shinohara, Franz Klein, Susan M. Gasser, Akira Shinohara

**Methodology:** Kiran Challa, Miki Shinohara, Susan M. Gasser

**Project administrations:** Akira Shinohara

**Resources:** Kiran Challa, Miki Shinohara

**Supervision:** Akira Shinohara

**Validation:** Kiran Challa

**Visualization:** Kiran Challa, Ghanim Fajish V, Akira Shinohara

**Writing - original draft:** Akira Shinohara

**Writing – review & editing:** Kiran Challa, Miki Shinohara, Franz Klein, Susan M. Gasser, Akira Shinohara

