## Supplementary figures and images for "Meiosis-specific prophase-like pathway controls cleavage-independent release of cohesin by Wapl phosphorylation"

### Supplementary file 1

#### Challa et al. Figure S1

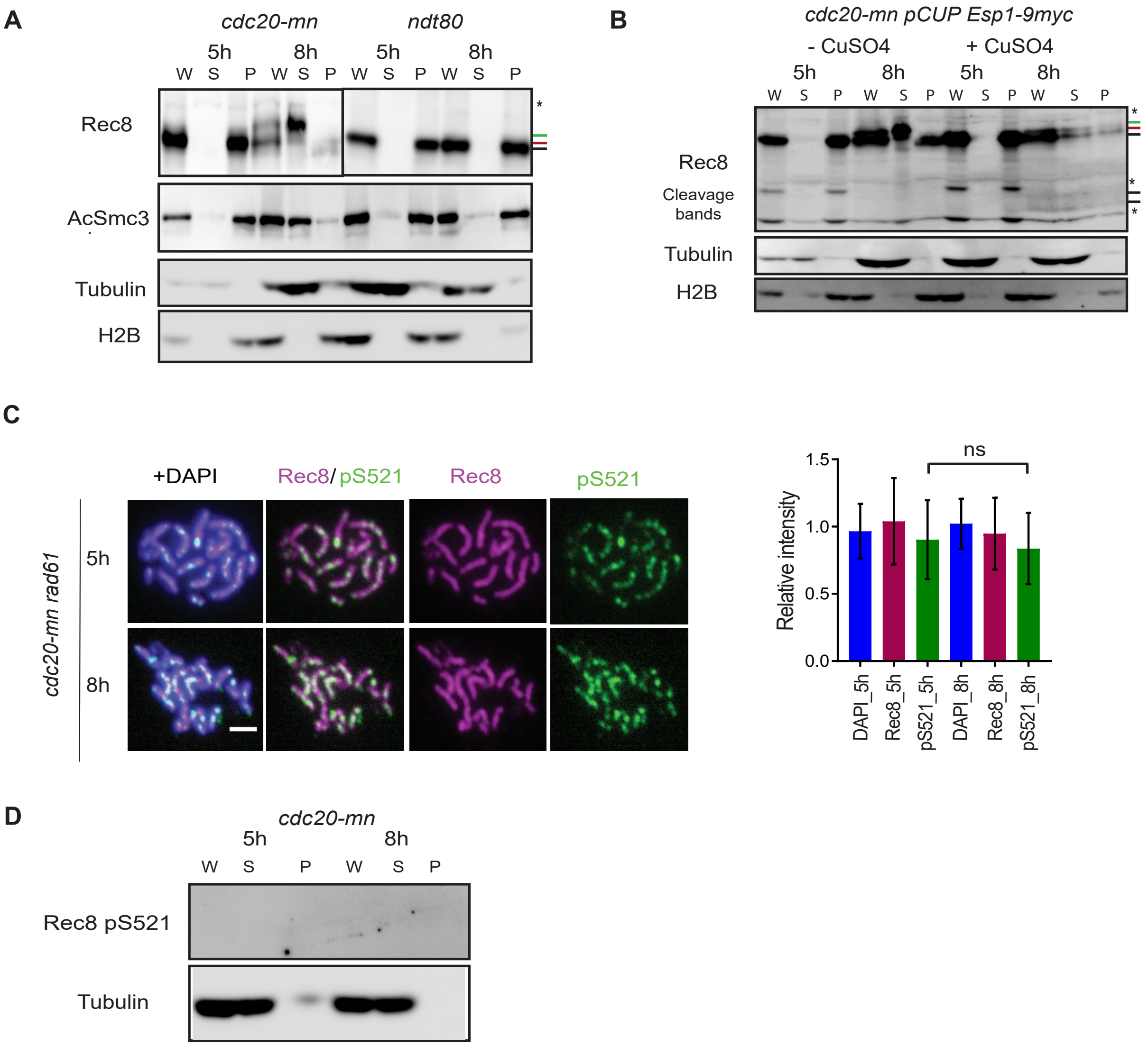

Challa et al. Figure S2

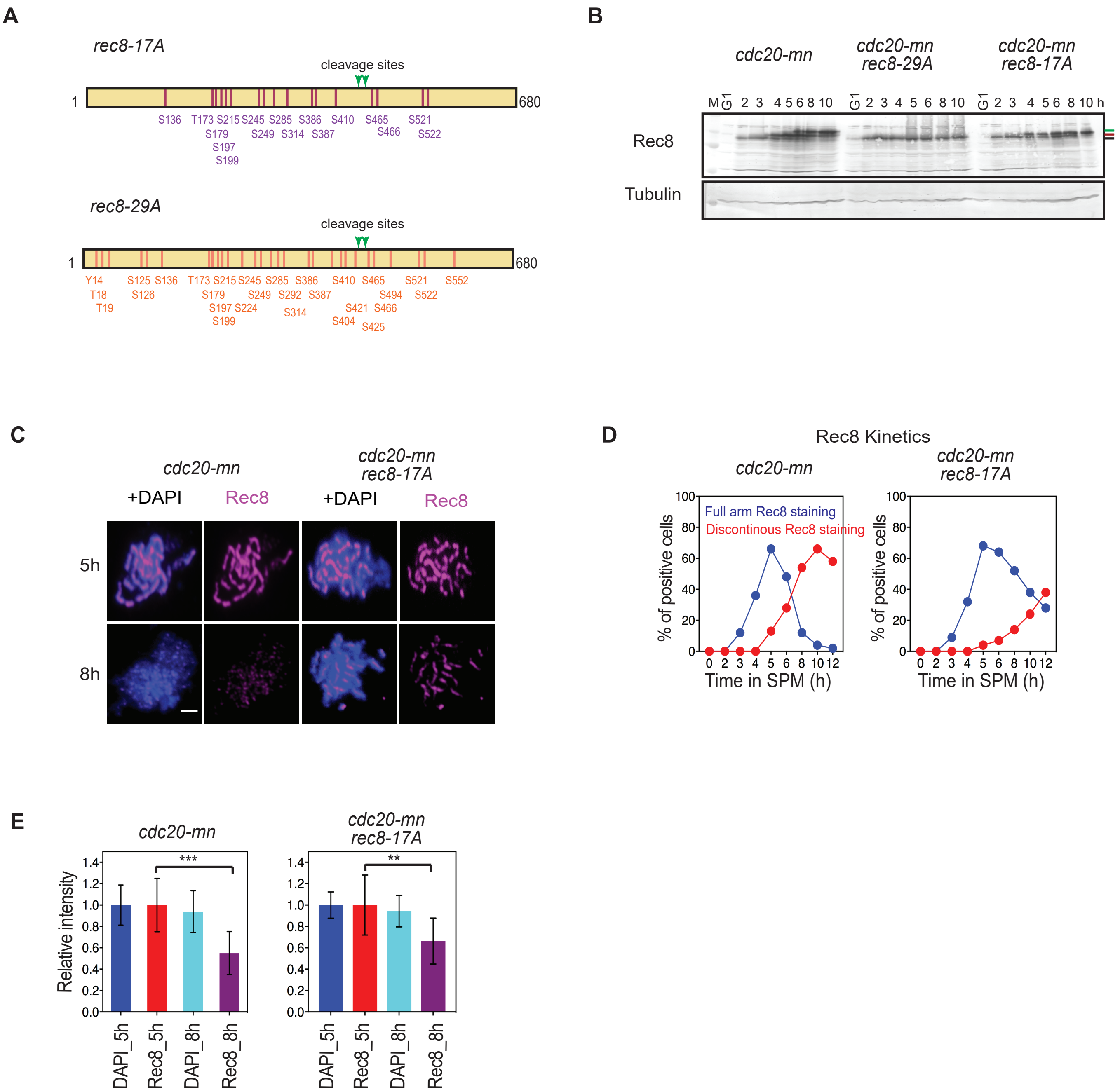

#### Challa et al. Figure S3

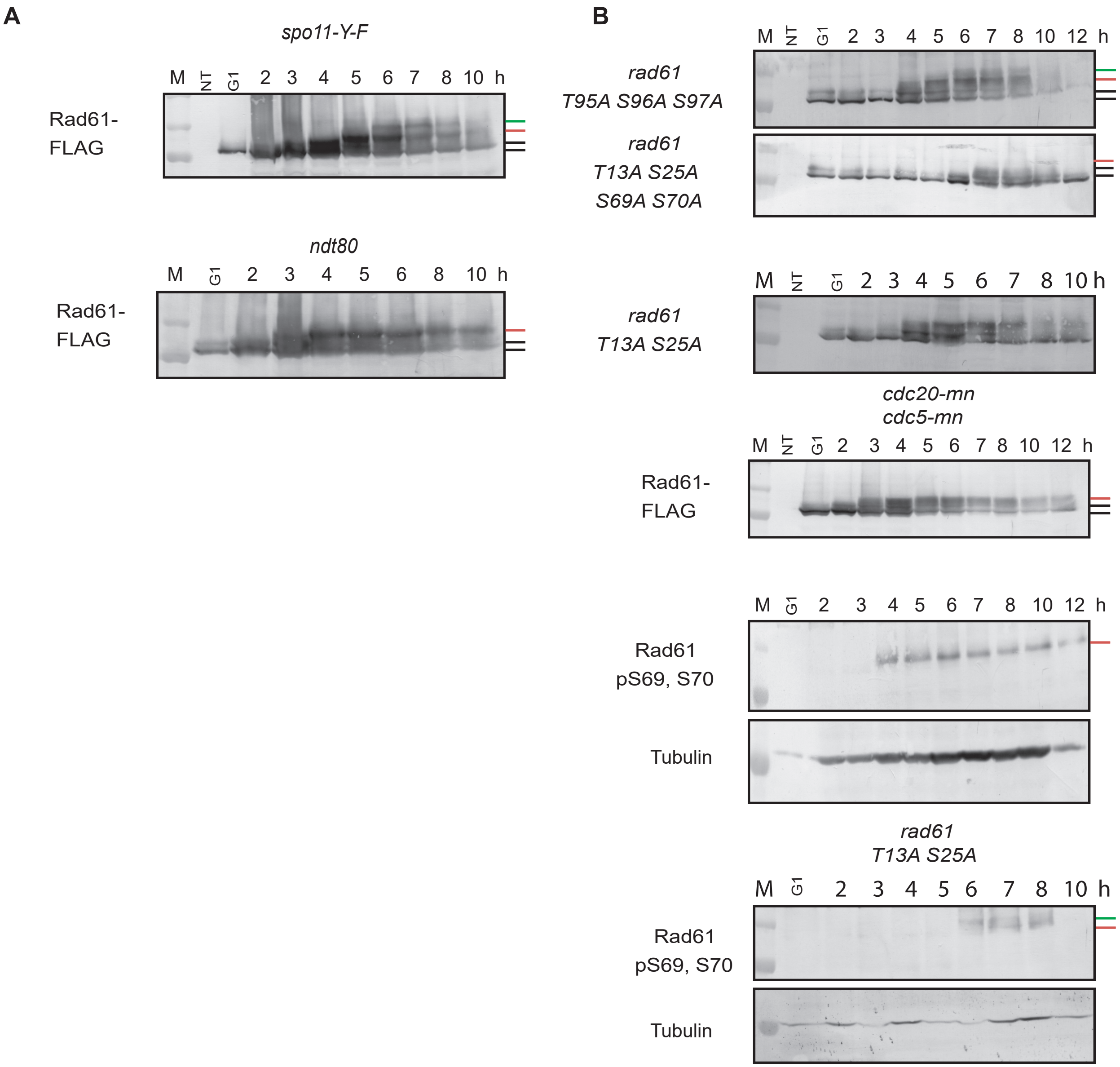

##### Challa et al. Figure S4

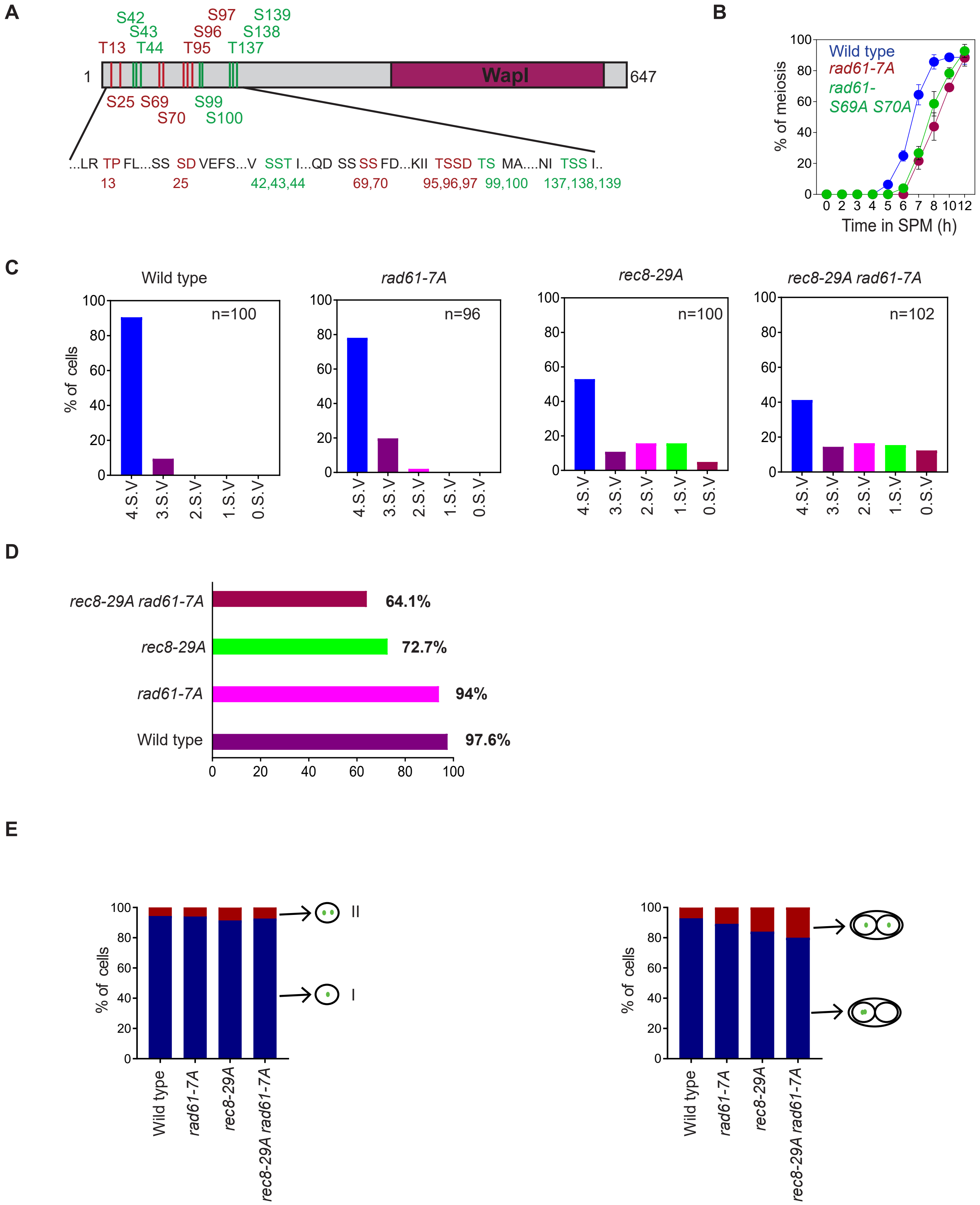
